# The relationship between land cover and microbial community composition in European lakes

**DOI:** 10.1101/2021.11.17.468820

**Authors:** Theodor Sperlea, Jan Philip Schenk, Hagen Dreßler, Daniela Beisser, Georges Hattab, Jens Boenigk, Dominik Heider

**Author notes:** Corresponding author: Dominik Heider.

## Abstract

Microbes such as bacteria, archaea, and protists are essential for element cycling and ecosystem functioning, but many questions central to the understanding of the role of microbes in ecology are still open. Here, we analyze the relationship between lake microbiomes and the land cover surrounding the lakes. By applying machine learning methods, we quantify the covariance between land cover categories and the microbial community composition recorded in the largest amplicon sequencing dataset of European lakes available to date. We identify microbial bioindicators for these land cover categories. Combining land cover and physico-chemical bioindicators identified from the same amplicon sequencing dataset, we develop two novel similarity metrics that facilitate insights into the ecology of the lake microbiome. We show that the bioindicator network, i.e., the graph linking OTUs indicative of the same environmental parameters, corresponds to microbial co-occurrence patterns. Taken together, we demonstrate the strength of machine learning approaches to identify correlations between microbial diversity and environmental factors, potentially opening new approaches to integrate environmental molecular diversity into monitoring and water quality assessments.

## 1 Introduction

Ecosystems are governed by processes at very different scales, ranging from the level of chemical reactions that shape the metabolism of single organisms to processes at the landscape level (Allen et al., 2014, Dobzhansky, 1966, Odum, 1997). Furthermore, the different scales are interlinked. For example, lakes and other freshwater habitats accumulate water from their catchment, and with it, nutrients, stressors, and pollutants. Because a lake’s water quality depends on its watershed’s ability to collect and purify water, lakes can be considered sentinels of environmental change of the landscape they are part of (O’Neill et al., 1997, Williamson et al., 2008). Microbes, in turn, play an essential role in the functioning and the stability of ecosystems and can be regarded as “first responders” to environmental change (Colwell, 1997, Docherty and Gutknecht, 2011, Webster et al., 2018). Indeed, because of their sensitivity to changes in the physico-chemical makeup of their environment, microorganisms are increasingly being used as bioindicators for ecosystem integrity in monitoring schemes (Birk et al., 2012, Cordier et al., 2018, Hering et al., 2010, Kermarrec et al., 2014).

While anthropogenic land cover change is one of the drivers of ecosystem quality decline (Intergovernmental Platform on Biodiversity and Ecosystem Services (IPBES), 2019, Song et al., 2018, Steinbauer et al., 2018), it is almost impossible to study its effects on freshwater ecosystems in well-controlled, experimental settings. Furthermore, the complex nature of ecosystems undermines the study of the interconnection between land cover and microbial community composition in field experiments and observational settings (Levin, 1998): The response of an organism to an environmental signal can be modulated by the presence and abundance of the other organisms in the ecosystem (Green and Sadedin, 2005, Levins and Lewontin, 1980, Wang et al., 2020). Because the interdependencies of environmental parameters induce confounding effects into our statistical analyses, they, furthermore, undermine our ability to detect direct and causal links. To make matters worse, in a fixed-size landscape, the areas of different land cover classes are not statistically independent from each other because an increase in one necessarily leads to decreases in others (King et al., 2005). Because of this, bioindicators (species identified by the indicator value function) can be used as apparent proxy measurement for the ecological variables they are indicative for (and therefore of use for biomonitoring schemes), but are not necessarily in a biologically relevant relationship with it (Landres et al., 1988, Simberloff, 1998).

While not being able to identify causal effects, with the required statistical and interpretative caution, insights into how freshwater microbiomes might be affected by land cover and land cover changes can still be gained. For example, recent studies have shown that microbial communities in waterways and lakes that are surrounded by different land cover types are significantly dissimilar (Kraemer et al., 2020, Saxena et al., 2015) and that land conversion for agricultural or urban uses influence the microbial communities of nearby stream sediments (Martin et al., 2020). Furthermore, it is well-known that the introgression of nitrates and other nutrients can lead to eutrophication and has a strong and characteristic effect on the microbial community composition (Gatti et al., 2018, Han et al., 2019, Sagova-Mareckova et al., 2021). Nevertheless, a systematic examination of the interrelation between land cover and microbial community composition for lakes in Europe is lacking for now.

In a prior publication, we developed the covariation framework, a statistical framework for the study of environmental microbiomes in observational settings such as environmental monitoring (Sperlea et al., 2021). The central idea behind this approach is to find a projection of the high-dimensional microbial community composition into the one-dimensional space of a target environmental parameter; calculating the *R*^2^ between this projection and the measured values of the environmental parameter in question corresponds to the amount of variation in the latter explained by the variation in the former and can be considered a measure of covariation. The covariation framework circumvents many of the obstacles described above: First, by estimating the covariation using machine learning methods that can model non-linear dependencies in non-independent data, like Random Forests (Breiman, 2001), as projection function, it handles the interdependencies between microbial species in the dataset. Second, it explicitly avoids any association with direct interaction, correlation, let alone causal relationships between the microbiome and the environmental parameters in question.

In this paper, we use the covariation framework as well as other machine learning-based approaches to study the relationship between land cover surrounding a lake and the microbiome of the lake. To this end, we analyze the largest amplicon sequencing dataset of European lakes available to date in concert with land cover data from the OpenStreetMap (OSM) project as well as the CORINE Land Cover (CLC) dataset from the Copernicus Land Monitoring Service (European Union, 2012, OpenStreetMap contributors, 2017). The former of the two data sources provides an open, community-driven, and, thus, rather detailed but potentially incomplete land cover categorization. In contrast, the latter dataset is based on high-resolution satellite imagery and contains a hierarchical categorization of land cover in 44 classes. Based on these analyses, we identify multi-target bioindicators, i.e., species indicative of multiple environmental parameters, as well as environmental parameters that might act as drivers of microbial community composition. Furthermore, we propose a novel data abstraction, the response map, that clusters environmental parameters in terms of the response they engender in the microbiome. Aside from providing targets for future experimental investigation, the results presented here highlight that aggregation of variables when studying ecosystems can obscure real relationships. Finally, this study provides a thorough analysis of the response of the lake microbiome with regard to a wide range of land cover and physico-chemical parameters.

## 2 Materials and methods

### 2.1 Amplicon sequencing

Sampling was part of a pan-European study conducted in August 2012 (eukaryotic sequences are published in (Boenigk et al., 2018); NCBI Bioproject PRJNA414052, prokaryotic sequences are published and described in (Nuy et al., 2020) and (Bock et al., 2020); NCBI Bioproject PRJNA559862). Methods for data collection, extraction, sequencing, and amplicon processing are described in detail in these studies (Bock et al., 2020, Boenigk et al., 2018, Nuy et al., 2020) and will be briefly outlined below.

To analyze bacterial and protistan freshwater communities on a large scale, 280 lakes were sampled throughout Europe. Sampling details and information on measured physico-chemical and geographical parameters can be found in (Boenigk et al., 2018). For DNA analyses, filtered water samples were air-dried and frozen in liquid nitrogen. Genomic DNA was extracted using the my-Budget DNA Mini Kit (Bio-Budget Technologies GmbH, Krefeld, Germany) with modifications after (Boenigk et al., 2018). Amplicon sequencing targeted the V2-V3 region of the 16S rRNA gene for bacteria, the V9 region of the 18S, and the ITS1 gene for eukaryotes. Samples were commercially sequenced (Fasteris, Geneva, Switzerland) on an Illumina HiSeq 2500 sequencer generating 300 bp long paired-end reads. Adapter removal, quality trimming, and demultiplexing were performed by the sequencing company.

Sequence processing was performed using a provisional version of the Natrix pipeline (Welzel et al., 2020). If not stated otherwise, all software versions and parameters were used as described in (Welzel et al., 2020). The main steps included quality checks using FASTQC (Andrews, 2010) and PRINSEQ (Schmieder and Edwards, 2011), assembly of paired-end reads with PANDASeq (Masella et al., 2012) and dereplication and chimera removal using UCHIME (usearch v7.0.1090 with default parameters) (Edgar et al., 2011). AmpliconDuo (Lange et al., 2015) was used to discard sequences that were not found in both technical replicates. The remaining sequences were clustered using SWARM (Mahé et al., 2014) and further aggregated to identical V9 sequences. This aggregation served as the basis for the OTU tables. The taxonomic assignment of the eukaryotic sequences was performed by a BLAST search (Altschul et al., 1990) against the NCBI nt database (from Dec 5, 2017) and for the prokaryotic sequences against SILVA SSURef 132 (Quast et al., 2012). For all downward analyses, we combined the prokaryotic and eukaryotic OTU tables.

### 2.2 Land cover data

Two different land cover datasets were used in this study. For both, we accessed data for the year 2012 because this was also the year the lake samples were collected. The CLC dataset was downloaded from the official website of Copernicus Earth Observation program (CLC 20212, v.2020 20u1, 100m raster GeoTiff)(European Union, 2012). The relative areas of the land cover classes were extracted from the dataset for circular areas around the sampling points with different radii using QGIS 3.16 (QGIS Development Team, 2020). Areas were aggregated to higher-level land cover classes according to the hierarchical CLC class model.

OSM land cover data was extracted from the OSM planet file from September 2012 archived at archive.org. This file was loaded in a PostgreSQL database and queried using a routine adapted from SEDE-GPS (Sperlea et al., 2018) to retrieve the map tiles surrounding the sampling position, to fuse these, and to extract a circular area of a given radius. Map tiles were rendered using the default mapnik map style, which was adjusted to (i) merge pixels of land use sub-categories with the respective main category (such as “tertiary road” with “road”) and (ii) remove signs, labels, and point of interest markers. The pixel-areas summarised per unique category of the resulting image were read out and translated back to meters.

Outlier land cover values in all subsets (concerning both the radius as well as the land cover category) of these two datasets were detected using the function *boxplot.stats* in R 4.0.3 for the different radii and land cover categories, separately. Samples containing an outlier or a value of zero for a given land cover category at the given radius were discarded for the analysis of the respective land cover category and radius.

Additional physico-chemical parameters were taken from (Sperlea et al., 2021).

### 2.3 Microbial biodiversity and land cover

A straightforward way of determining whether land cover changes impact lake microbiomes is to assess whether the distribution of land cover types surrounding the lake is predictive of the lake’s microbial biodiversity. To this end, we extracted the relative area covered by different land cover categories in circular areas around the sampling sites of the European lake dataset from both the CORINE land cover (CLC) dataset as well as the OpenStreetMap (OSM) project (see Methods). These two datasets differ in the way they were generated and their categorization of land cover. While the former is derived from satellite data, the latter is annotated in a community-driven manner, based on landscape features observed “on the ground”. To distinguish between effects present at shorter or longer geographic ranges, we extracted and analyzed areas surrounding the sampling points within radii ranging from 1 km to 10 km, in steps of 1 km, as well as 25 km and 50 km. We then assessed the degree to which the relative sub-areas of the land cover categories contained in the extracted areas can be used to predict a set of biodiversity metrics calculated for the microbial communities of the sampled lakes using Random Forest models.

To calculate biodiversity metrics, the OTU table was rarefied using the *rrarefy* function from the R package `vegan` (v2.5-6, ref. Oksanen et al., 2019). Biodiversity metrics were calculated from rarefied OTU tables using the *diversity* (Shannon index, Simpson diversity, inverse Simpson diversity), and *renyi* (Renyi entropy) functions from the R package vegan, except for species richness, which is the total number of OTUs present, and Pielou’s evenness, which was calculated by dividing the sample’s Shannon index by the log of the sample’s richness (Chao and Jost, 2015, Daly et al., 2018). Renyi entropy metrics were calculated for *α* = {0.25, 0.5, 1, 2, 4, 8, 16, 32, 64, ∞}, as different values for this parameter drastically change the metrics sensitivity to relative species abundance (Chao and Jost, 2015). For the prediction of biodiversity metrics based on land cover categories, Random Forests from the R package `caret` (version 6.0.86, ref. Kuhn, 2008) were trained using 10-fold cross-validation without further feature selection.

### 2.4 Covariation framework

Covariation between the lake microbiomes and land cover areas was quantified using the covariation framework presented in ref. Sperlea et al., 2021. Methodologically, the framework is a straightforward machine learning approach, training a machine learning model to predict the values of an environmental parameter after feature selection. However, the model’s prediction is interpreted as the projection of the microbial community composition to the space of the target variable while leveraging the model’s potential to model non-linear interdependencies in the microbiome. This way, the coefficient of determination, *R*^2^, can be interpreted as a measure of covariation between the microbiome as a whole and the target variable. As feature selection method, the *multipatt* function (with the parameter “indval”) from the `indicspecies` R package (v1.7.9, ref. Cáceres and Legendre, 2009) was used after Hellinger transformation (Legendre and Gallagher, 2001) of the OTU counts to identify bioindicator OTUs for tertiles of the respective land cover category. Random Forest models from the R package `caret` (version 6.0.86, ref. Kuhn, 2008) were trained in a 10-fold cross-validation scheme with the OTU tables as independent and the relative area of a single land cover category as dependent variables, with both being centered and log-ratio transformed. Because of statistical limitations of the Random Forest model, some combinations of area size and land cover category could not be used for model training.

Confidence intervals for the model evaluations were estimated based on resampling of predicted and measured dependent variable pairs with replacement with thousand repetitions. Statistical significance of relevant models was asserted by comparing the R^2^ value with results gathered by thousand repetitions of training models with the same hyper-parameter setting on resampled biodiversity data in a Student’s t-test as implemented in the *t.test* function in R 4.0.3.

### 2.5 Bioindicator analysis

Bioindicator OTUs for land cover categories were identified using the indicator species method as implemented in the *multipatt* function in the `indicspecies` R package (v1.7.9) Cáceres and Legendre (2009), Dufrêne and Legendre (1997). A significance level of *α* = 0.05 was applied after Benjamini-Hochberg correction for the total number of land cover categories analyzed in this study.

The similarity of two environmental parameters was calculated in terms of the microbiome’s response to changes in them by calculating the Jaccard similarity between the lists of OTUs indicative of the two parameters. The resulting similarity matrix was visualized as a force embedded network using the function *qgraph* from the package `qgraph` (v1.6.9, ref. Epskamp et al., 2012). Furthermore, a dissimilarity matrix was derived from the similarity matrix by inversion after the addition of a random number in the order of 10^−8^ in order to avoid the division by zero. This dissimilarity matrix was visualized as a dendrogram using *upgma* and *ggdendrogram* from the packages `phangorn` (v2.5.5., ref. Schliep, 2011) and `ggdendro` (v0.1.22, ref. de Vries and Ripley, 2020), respectively. For the ordination of the environmental parameters, the *metaMDS* function from the R package `vegan` was used.

For the bioindicator network, an edge was created between all pairs of bioindicator OTUs that are indicative of at least one common environmental parameter. Null-hypothesis networks were created the same way based on resampled indicator lists; for these, each lake parameter is assigned the same number of randomly selected OTUs as pseudo-indicators in such a way that the distribution of cardinalities is the same as that of the real OTUs. Node properties (degree, closeness centrality, eigenvector centrality, page rank, and authority score) were calculated using the `igraph` package in R 4.0.3 (v1.2.6, ref. Csardi and Nepusz, 2006).

### 2.6 Network inference methods

In this study, we apply several methods for the inference of network structures from OTU tables. Most of these employ similarity measures as edge weights calculated between all pairs of OTUs, which act as nodes in the network. Simple co-occurrence, checkerboard score (Connor and Simberloff, 1979, Stone and Roberts, 1990), Bray-Curtis similarity (Bray and Curtis, 1957), Kullback-Leibler divergence (Kullback and Leibler, 1951), Pearson and Spearman correlation were used as similarity metrics. The co-occurrence metric was defined as the number of samples in which the two OTUs in question had non-zero occurrence. Pearson and Spearman correlations were calculated using the *cor* function in R and results below the significance level *α* = 0.05 were discarded. For the calculation of the Kullback-Leibler divergence, zeroes in the OTU table were replaced by 10^−8^ to avoid infinities created by the logarithm. Additionally, the method SparCC (Friedman and Alm, 2012) was used as implemented in the *sparcc* function from the `SpiecEasi` package (v1.1.0, ref. Kurtz et al., 2015) with default parameters. For all networks, OTUs that have zero counts for 25 or more sampling sites were excluded from the analysis to avoid statistical artifacts that are based on the rarity of the OTUs in question (Berry and Widder, 2014).

All figures were generated using the R packages `ggplot2` (v.3.3.2, ref. Wickham, 2016), unless otherwise noted, and following the guidelines laid out in ref. Hattab et al., 2020.

## 3 Results

### 3.1 Microbial biodiversity in lakes only marginally reflects differences in surrounding land cover

A straightforward way of determining whether land cover changes impact lake microbiomes is to assess whether the distribution of land cover types surrounding the lake is predictive of the lake’s microbial biodiversity. To this end, we extracted the relative area covered by different land cover categories in circular areas around the sampling sites of the European lake dataset from both the CORINE land cover (CLC) dataset as well as the OpenStreetMap (OSM) project (see Methods). To be able to distinguish between effects present at shorter or longer geographic ranges, we extracted and analyzed areas surrounding the sampling points within radii ranging from 1 km to 10 km, in steps of 1 km, as well as 25 km and 50 km. We then assessed the degree to which the variation in the relative sub-areas of the land cover categories contained in the extracted areas explain the variation in microbial alpha diversity of the sampled lakes using Random Forest models. A set of alpha diversity metrics was used instead of a single metric because they capture different mathematical characteristics of biodiversity (Daly et al., 2018).

Our results show that there is, at best, a marginal relationship between land cover and microbial biodiversity, as no combination of radius, biodiversity metric, and dataset results in an *R*^2^ > 0.2 (see figure 1A, supplementary figure 1 and 2). For all radii studied here, the lake microbiome’s Renyi entropy is most predictable from land cover, followed by species richness. Furthermore, we found no significant difference between *R*^2^ values obtained for the same biodiversity metric at different radii (see figure 1B), indicating that the results presented here are most likely due to statistical artifacts rather than processes that shape microbial biodiversity based on surrounding land cover. In general, the land cover data collected from the OSM dataset is less predictive for microbial biodiversity than the CLC datasets (see figures 1A and B). Therefore, we focus our further analysis on the CLC dataset. Taken together, these results suggest that if land cover has a structuring effect on lake microbiomes, this relationship is not reflected in alpha diversity metrics.

**Figure 1:**
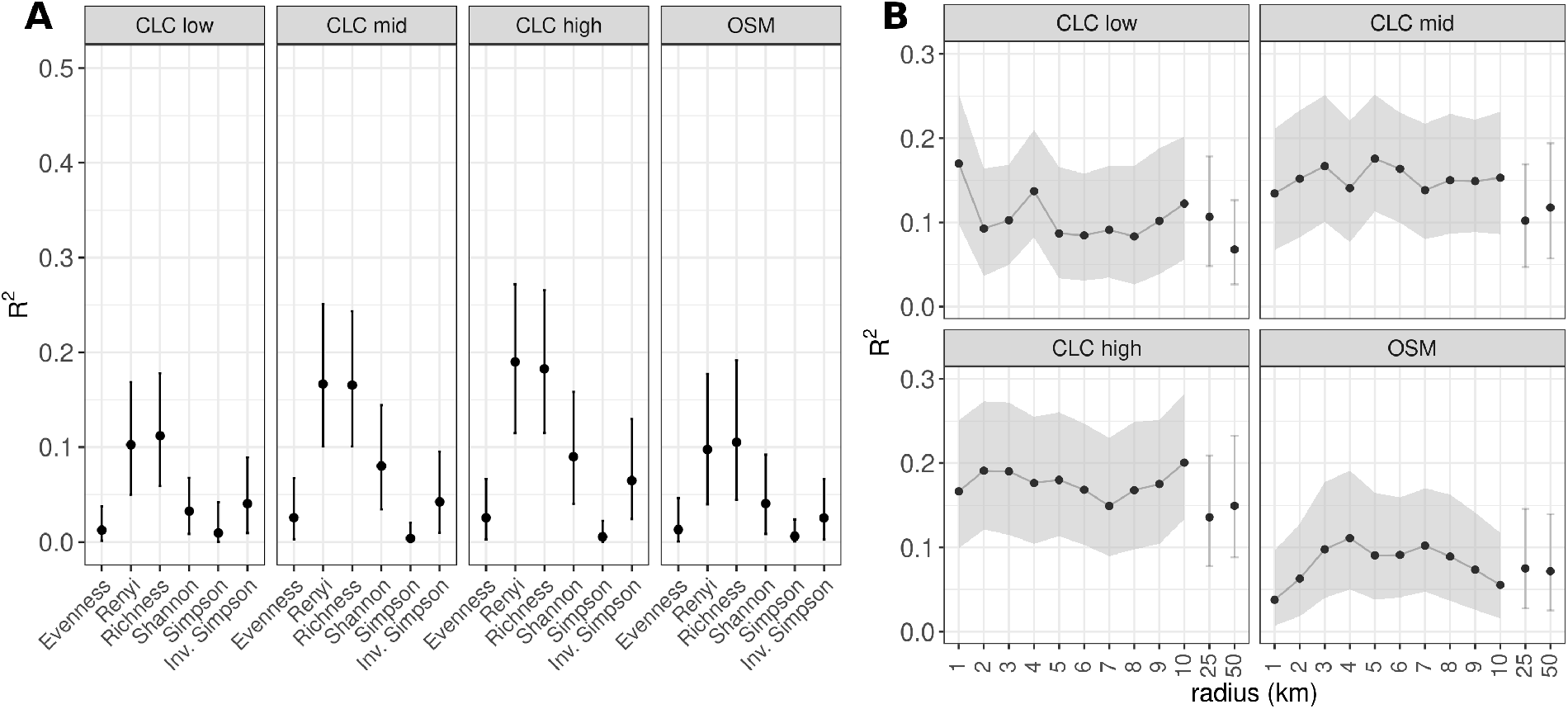
Evaluation of Random Forest models trained to predict biodiversity metrics from land cover. **A.** Results for the land cover in a radius of 3 km around the sampling site and all biodiversity metrics surveyed in this study. Lines represent confidence intervals estimated based on resampling (see methods). For results of other radii, and results for Renyi entropy with other values for *α*, see supplementary figures 1 and 2. **B.** Results for the full range of radii as well as 25 and 50 km for Renyi entropy with *α* = 0.5. Grey areas and error bars represent confidence intervals; results for 25 and 50 km are visually separated to underline that these radii are not in the range of the other radii.

### 3.2 Microbial community structure covaries with specific land cover categories

While not visible at the level of alpha diversity, we hypothesized that there must be an apparent relationship between land cover surrounding the lakes and the lake microbiome when analysing it at the fine-grained level of microbial community composition. To investigate this hypothesis, we applied the covariation framework to the OTU tables and the relative areas of the land cover categories present in the CLC datasets. Higher *R*^2^ values indicate a higher degree of covariation between the microbiome as a whole and the target parameter, i.e., the relative area of the land use category in question. On level 1 of CLC category hierarchy, we observe covariation of *R*^2^ > 0.05 for “artificial surfaces (1)” and “agricultural areas (2)” at very low radii as well as increasing covariation for “forest and semi-natural areas (3)” with increasing radii (see figure 2A). In contrast, the anthropogenic effects that are expected with built surfaces and agriculturally used land (Gatti et al., 2018, Martin et al., 2020) act on rather short ranges.

**Figure 2:**
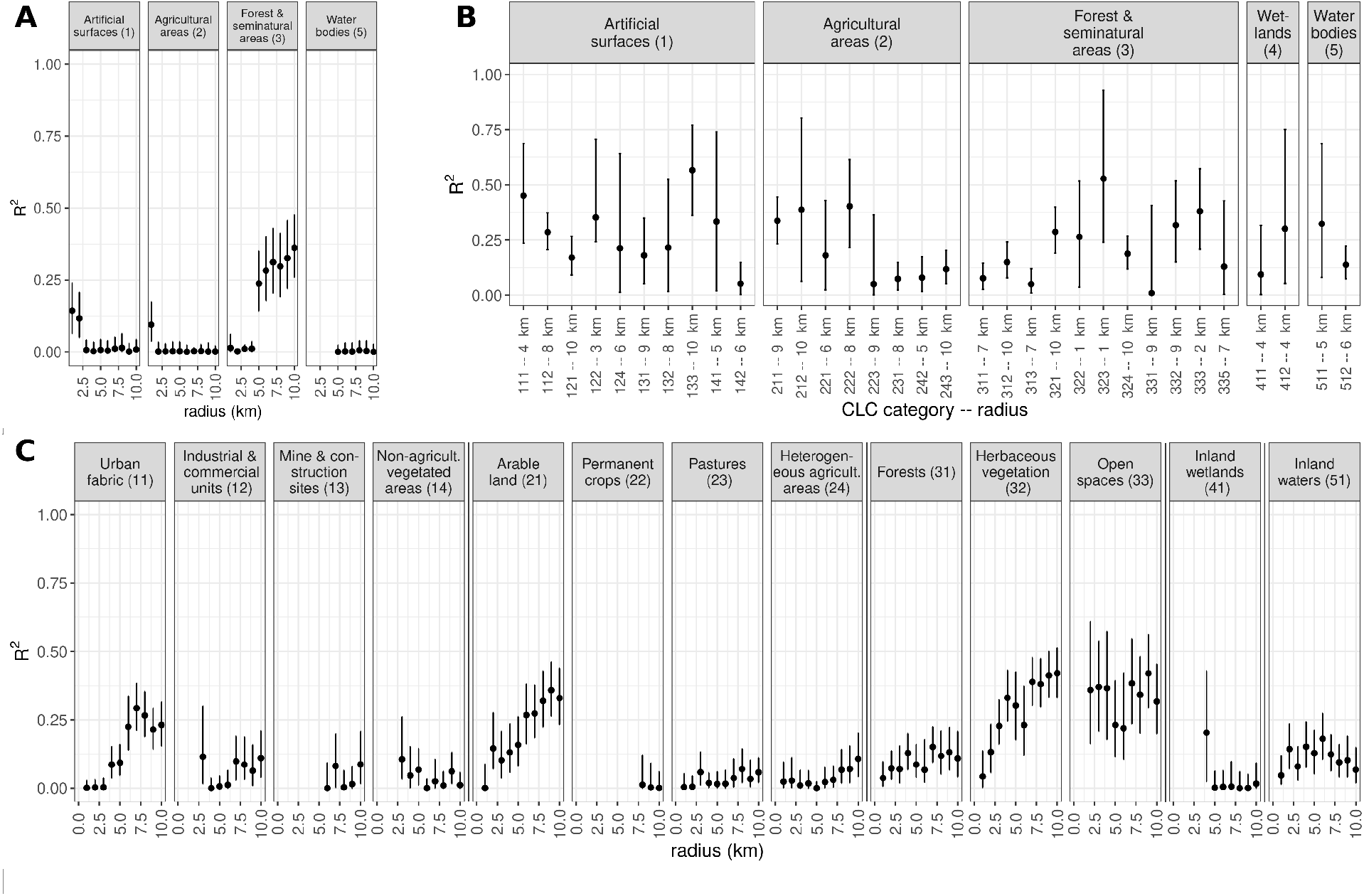
Covariation between land cover surrounding lakes and the lake’s microbiome. Numbers in brackets (**A** and **C**) and in x-axis labels (**B**) refer to the CLC category number code (see table 1). For ease of display, full-length labels of the CLC categories have been shortened in some cases. Vertical lines represent confidence intervals estimated from resampling (see methods) and dots represent covariation as obtained in model evaluation. **A.** Results for all high-level land cover categories in the CLC dataset. **B.** Results for low-level land cover categories; for each land cover category, only the results for the radius with the highest *R*^2^ are shown. **C.** Results for all mid-level land cover categories in the CLC dataset. Vertical lines between facets separate groups of categories that are subcategories of the categories in **A**. For all results, see supplementary table 1.

The covariation observed between the lake microbiome and land cover categories at level 2 of the CLC hierarchy paints a more nuanced picture (see figure 2C). For example, we observe increasing covariation with the lake microbiomes at increasing radii for the land cover categories “arable land (21)” and “scrub and/or herbaceous vegetation associations (32)”. In contrast, for “urban fabric (11)” and “inland waters (51)”, we observe a peak in covariation at radii of 7 km and 6 km, respectively, with lower *R*^2^ for the other radii. For “forests (31)” and “open spaces with little or no vegetation (33)”, the covariation for different radii stays within the respective confidence intervals of the covariation at the 1 km radius. Notably, the covariation between the microbiome and sub-categories of a CLC category often deviate strongly from the covariation between the microbiome and the respective super-category. The same applies to covariations observed at level 3 of the CLC category hierarchy (see figure 2B). Taken together, these results show that a broad array of land cover categories have an impact on the lake’s microbial community composition at the OTU level and they do so at different radii. This can be interpreted as a reflection of different mechanisms being at play for the influence of lake ecology of different land cover categories. Furthermore, the more specific land cover categories at levels 2 and 3 of the CLC category hierarchy generally show higher levels of covariation with the lake microbiome, indicating that aggregation of land cover classes can hide relationships apparent at more fine-grained levels of analysis.

To identify general spatial trends, we separately calculated the mean covariation of all land cover categories for each radius and land cover hierarchy level. For all but one radius-hierarchy level combination, the average *R*^2^ value is below 0.15 (see supplementary figure 3) and throughout all combinations, the relative standard deviation is close to or higher than 100%. This shows that there are neither general spatial trends nor a generally higher covariation at higher levels of the CLC hierarchy. Instead, to observe specific, radius-dependent effects of land cover on the lake microbiome, it is important to differentiate between land cover categories.

### 3.3 Microbial lake bioindicators for surrounding land cover categories

Using the indicator value method that is part of the covariation framework, we identified 2,354 OTUs that act as bioindicators for the land cover categories in a total of 4,453 indicator-parameter pairs (for a complete list, see supplementary table 2). Among the land cover categories studied in this paper, for “scrub and/or herbaceous vegetation associations (32)” and “forest and semi-natural areas (3)” we identify the highest number of indicator OTUs with 1056 and 703 OTUs, respectively (see table 1). Most of the indicator OTUs are Bacteria (87%) from the phyla Proteobacteria (29%), in particular Alphaproteobacteria (13%), Bacteroidetes (28%), in particular Flavobacteriia (16%), or Cyanobacteria (15%). Furthermore, most of the OTUs obtained are indicative for more than one land cover parameter (fig. 3A). All OTUs indicative for more than seven land cover parameters are bacteria (see tables 2 and 3). Taken together, these results support the notion that bacteria are more sensitive to environmental changes or respond to environmental signals in a different manner than microbial eukaryotes (Bock et al., 2020, Logares et al., 2018).

**Figure 3:**
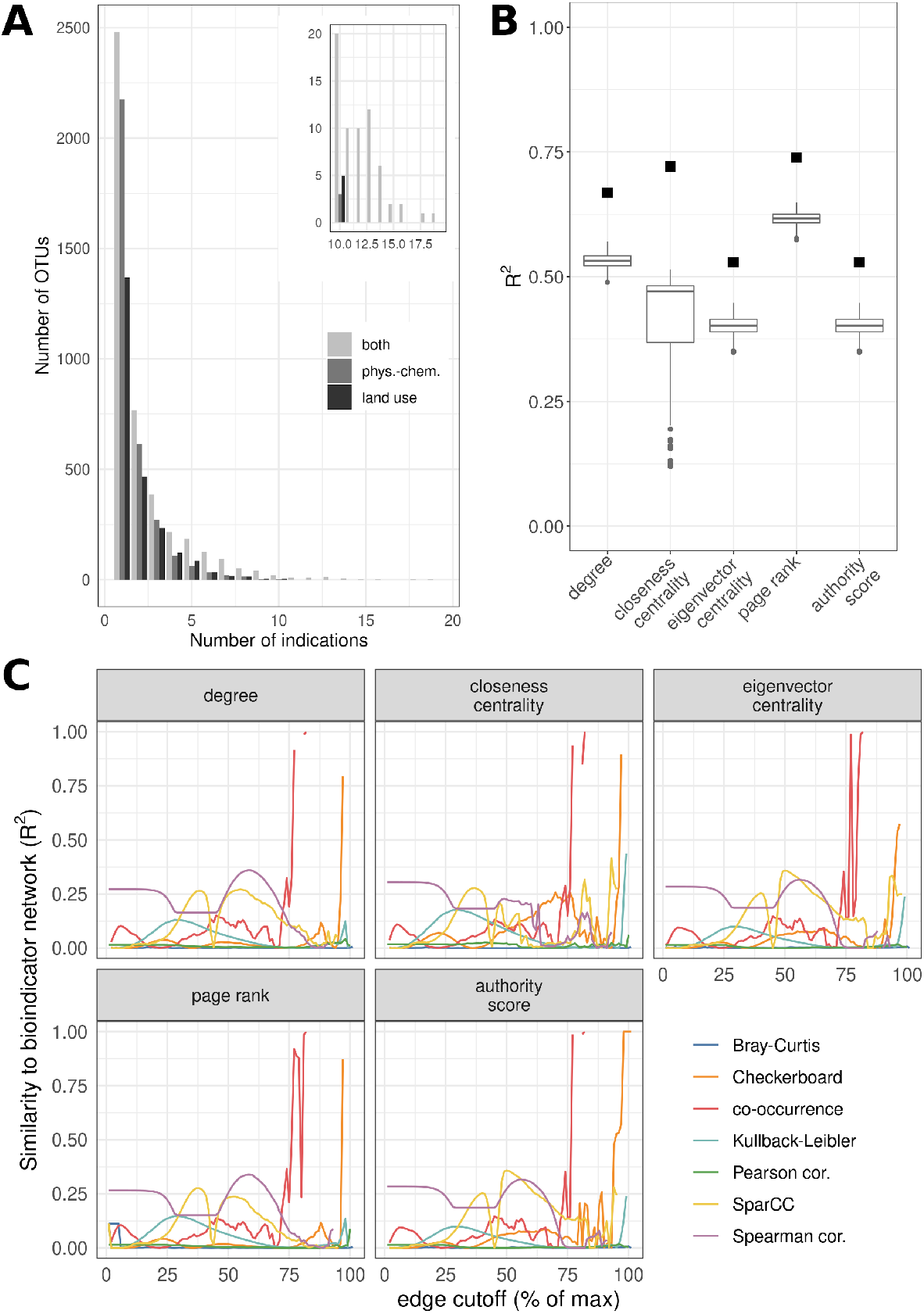
The node properties of the bioindicator network are significant and comparable to co-occurrence networks. **A.** Distribution of multitask bioindicators across numbers of indicated land cover categories. Inset: Distribution of multitask bioindicators for ten or more parameters. **B.** Correlation between the cardinality of each bioindicator OTU and the node properties of the respective node in the bioindicator network. Black squares: Results for the bioindicator network. Grey box plots: Results for resampled null-hypothesis networks (for details, see methods). For all metrics, the results for the bioindicator network are significantly different from those of the null-hypothesis networks (one-sample t-test, P< 2.2e 16). **C.** Comparison of the bioindicator network and established methods for the network inference from OTU tables. Networks are compared by the coefficient of determination, *R*^2^, between a property of the nodes in the bioindicator network and the network created using a network inference method. As some of the methods create fully connected networks by default, edges with weights smaller than a cut-off were removed; this cut-off ranges between the respective minimum and maximum edge weight in 100 equidistant steps.

**Table 1:** Numbers of microbial indicators for land cover categories (with respective radius, left) and physico-chemical parameters (right) and the respective *R*^2^ resulting from the covariation framework (results for physico-chemical parameters taken from Sperlea et al. (2021)).

| Parameter | Indicators | Radius (km) | $R^2$ | Parameter | Indicators | $R^2$ |
| --- | --- | --- | --- | --- | --- | --- |
| Herbaceous vegetation (32) | 1056 | 7 | 0.42 | Altitude | 1595 | 0.60 |
| Forest, semi-natural areas (3) | 703 | 10 | 0.36 | Conductivity (LF) | 1349 | 0.49 |
| Arable land (21) | 445 | 9 | 0.36 | Temperature | 920 | 0.54 |
| Non-irrigated arable land (211) | 416 | 9 | 0.34 | pH | 603 | 0.31 |
| Discontinuous urban fabric (112) | 276 | 8 | 0.29 | Sum of Ions | 166 | 0.50 |
| Agricultural areas (2) | 272 | - | - | Cations | 164 | 0.52 |
| Natural grasslands (321) | 265 | 10 | 0.29 | K<sup>+</sup> | 118 | 0.26 |
| Urban fabric (11) | 247 | 7 | 0.29 | Total Phosphorus (TP) | 92 | 0.31 |
| Artificial surfaces (1) | 177 | - | - | Anions | 89 | 0.53 |
| Forests (31) | 140 | 7 | 0.15 | Na<sup>+</sup> | 86 | 0.18 |
| Inland waters (51) | 77 | 6 | 0.18 | Mg<sup>2+</sup> | 70 | 0.34 |
| Industrial or commercial units (121) | 77 | 10 | 0.17 | DOC | 43 | 0.40 |
| Heterogeneous agricultural areas (24) | 69 | - | - | Ca<sup>2+</sup> | 32 | 0.37 |
| Transitional woodland-shrub (324) | 61 | 10 | 0.19 | HCO<sub>3</sub> | 27 | 0.40 |
| Open spaces with little vegetation (33) | 52 | 9 | 0.42 | NO<sub>3</sub> | 17 | 0.29 |
| Bare Rocks (332) | 34 | 9 | 0.32 | Alkalinity (Alk.Gran) | 16 | 0.38 |
| Water bodies (5) | 32 | - | - |  |  |  |
| Pastures (23) | 21 | - | - |  |  |  |
| Sparsely vegetated areas (333) | 15 | 2 | 0.38 |  |  |  |
| Permanent crops (22) | 6 | - | - |  |  |  |
| Fruit trees (222) | 3 | 8 | 0.40 |  |  |  |
| Inland wetlands (41) | 2 | 4 | 0.20 |  |  |  |
| Artificial, non-agricultural vegetated areas (14) | 2 | - | - |  |  |  |
| Industrial, commercial and transport units (12) | 2 | - | - |  |  |  |
| Vineyards (221) | 1 | 6 | 0.18 |  |  |  |
| Mineral extraction sites (131) | 1 | 9 | 0.18 |  |  |  |
| Mine, dump and construction sites (13) | 1 | 9 | 0.18 |  |  |  |
| Dump sites (132) | 1 | 8 | 0.22 |  |  |  |
| Construction sites (133) | 1 | 10 | 0.57 |  |  |  |

**Table 2:**
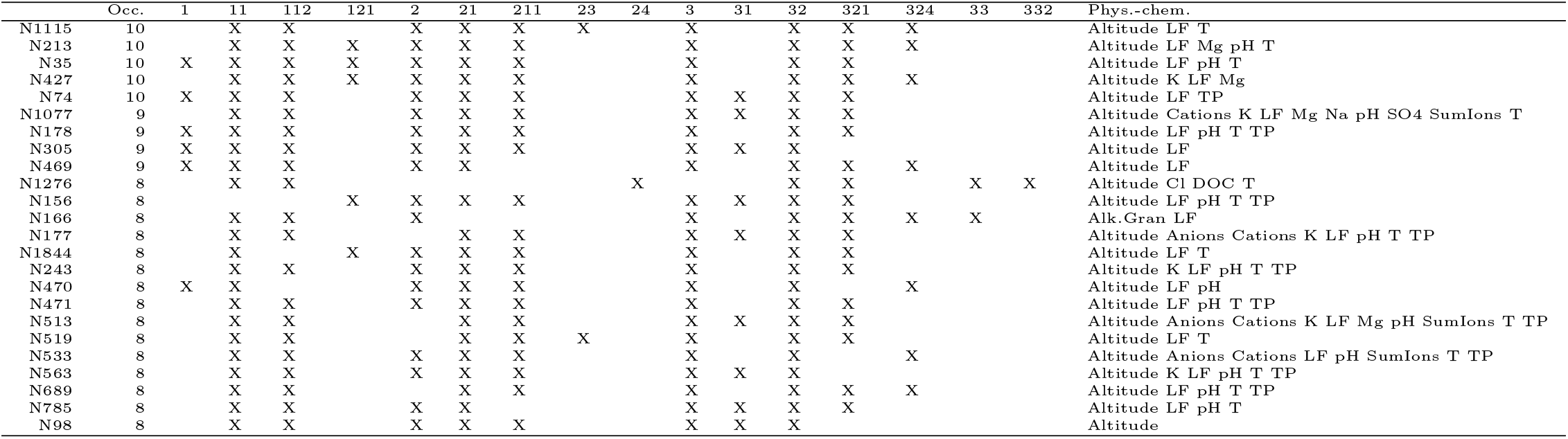
Multitask OTUs identified in this study for more than 6 land cover categories and the physico-chemical parameters they have been identified as indicators for in Sperlea et al. (2021). Numbers represent CLC land cover categories (see table 1). For taxonomic annotation of these OTUs, see table 3.

**Table 3:**
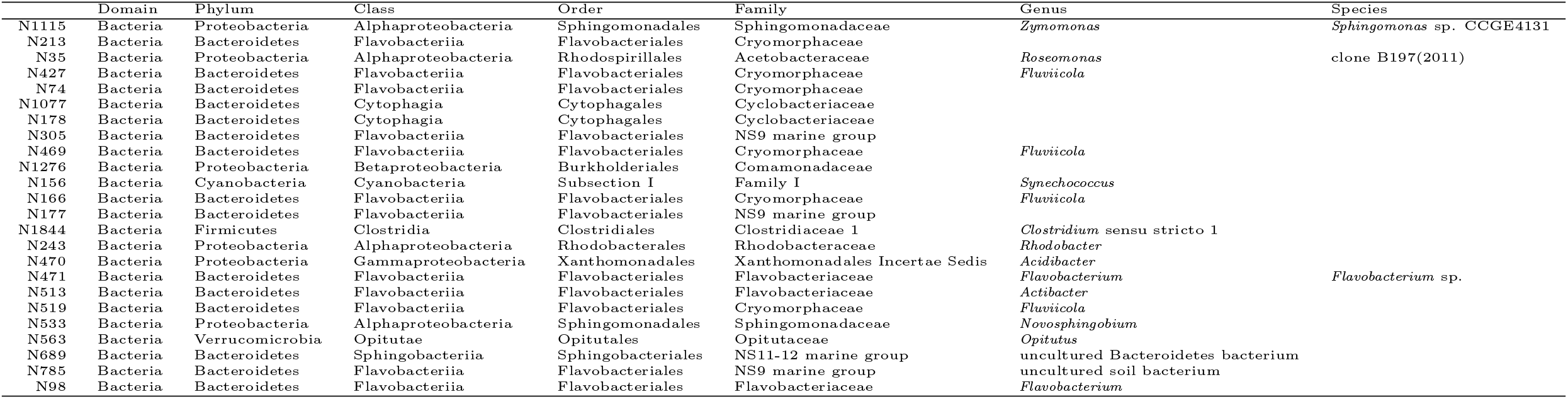
Taxonomic annotation of the multitask OTUs presented in table 2.

In a previous paper, we identified bioindicator OTUs for physico-chemical parameters while working with the same amplicon sequencing dataset as analyzed here Sperlea et al. (2021). Comparing the results of this analysis with those in the prior publication, we observed that almost all bioindicators indicative for at least eight land cover categories are also indicative of the lake’s altitude (see table 2). This underlines the central role the geographic location of a site plays in its ecology (see discussion) (Bock et al., 2020, Karlsson et al., 2005). However, a literature search for either the multi-target bioindicators on the Genus or Species level of taxonomy did not result in any information supporting or calling our results into question.

### 3.4 Structural insights into the lake microbiome from multitask bioindicators

To further elucidate the relationship between the lake microbiome and environmental parameters, we combined the bioindicator OTUs identified for land cover parameters with those for physico-chemical parameters identified in Sperlea et al. (2021). This way, we obtained a data structure that can be described as a set of maps between a set of OTUs and a set of environmental parameters. From this, we derived two distinct similarity matrices: One stating the similarity of OTUs in terms of the number of environmental parameters they are indicative of, and one that displays the similarity of the parameters in terms of the OTUs assigned to them.

The former can be turned into a bioindicator network as follows. Each bioindicator OTU is assigned to a node and edges are drawn between nodes representing OTUs that are indicative for at least one common environmental parameter (see supplementary table 3 for the entire network). We noticed correlations between the cardinality (i.e., the number of occurrences of an OTU in all bioindicator lists) of the nodes of the bioindicator network and the respective nodes’ degree, closeness centrality, eigenvector centrality, page rank, and authority score (see supplementary figure 4). Because this result could be due to basic graph properties, we compared the square of the Pearson correlation coefficient, *R*^2^, resulting from the correlation of cardinality with node properties of the bioindicator network with those gained from resampled networks (see Methods for details). We find that the nodes in the bioindicator network have statistically significant properties (see figure 3B), which suggests a biological relevance of the bioindicator network’s structure.

Furthermore, we compared the bioindicator network to networks inferred from the original OTU table. More specifically, we asked whether the node properties generated using network inference methods correlate with the node properties of the bioindicator network. We chose this approach to comparing the two network structures as it can capture relative differences between node properties and might thus be robust with regard to global effects of a network method, e.g., a consistently lower degree. Our results show that applying a high cut-off to co-occurrence and checkerboard score similarity matrices results in networks similar to the bioindicator network. In contrast, neither correlation-based nor compositionality-aware methods do so (see figure 3C).

The second data structure presents the similarity of pairs of environmental parameters in terms of the Jaccard similarity of the list of bioindicator OTUs identified for them. A high Jaccard similarity between two environmental parameters suggests that the microbiome responds to changes in the parameters in a similar manner. Along these lines, a visualization of this similarity matrix can be seen as a response map of the microbiome with regard to environmental changes. We attempted to visualize the resulting similarity matrix using non-metric dimensional scaling with up to 10 dimensions but were unable to receive stress values < 0.05, suggesting that the responses of the microbiome to environmental change are non-trivial. Nevertheless, the visualization of the similarity matrix as a UPGMA-derived dendrogram and an undirected graph (see figures 4A and B, respectively) results in multiple distinct clusters of highly similar environmental parameters. The largest one of these comprises the concentration of magnesium, potassium, anions, cations, phosphorus, as well as temperature, pH, altitude, and a wide range of land cover categories. Most of the clusters identified here can be explained with regard to physico-chemical and landscape ecological processes in lakes (see Discussion). A further notable result is the relatively high distances of many of the same CLC categories’ subcategories in both the dendrogram and the graph. This underscores our prior finding that lake microbiomes react to different land cover categories in different ways (see figure 2).

**Figure 4:**
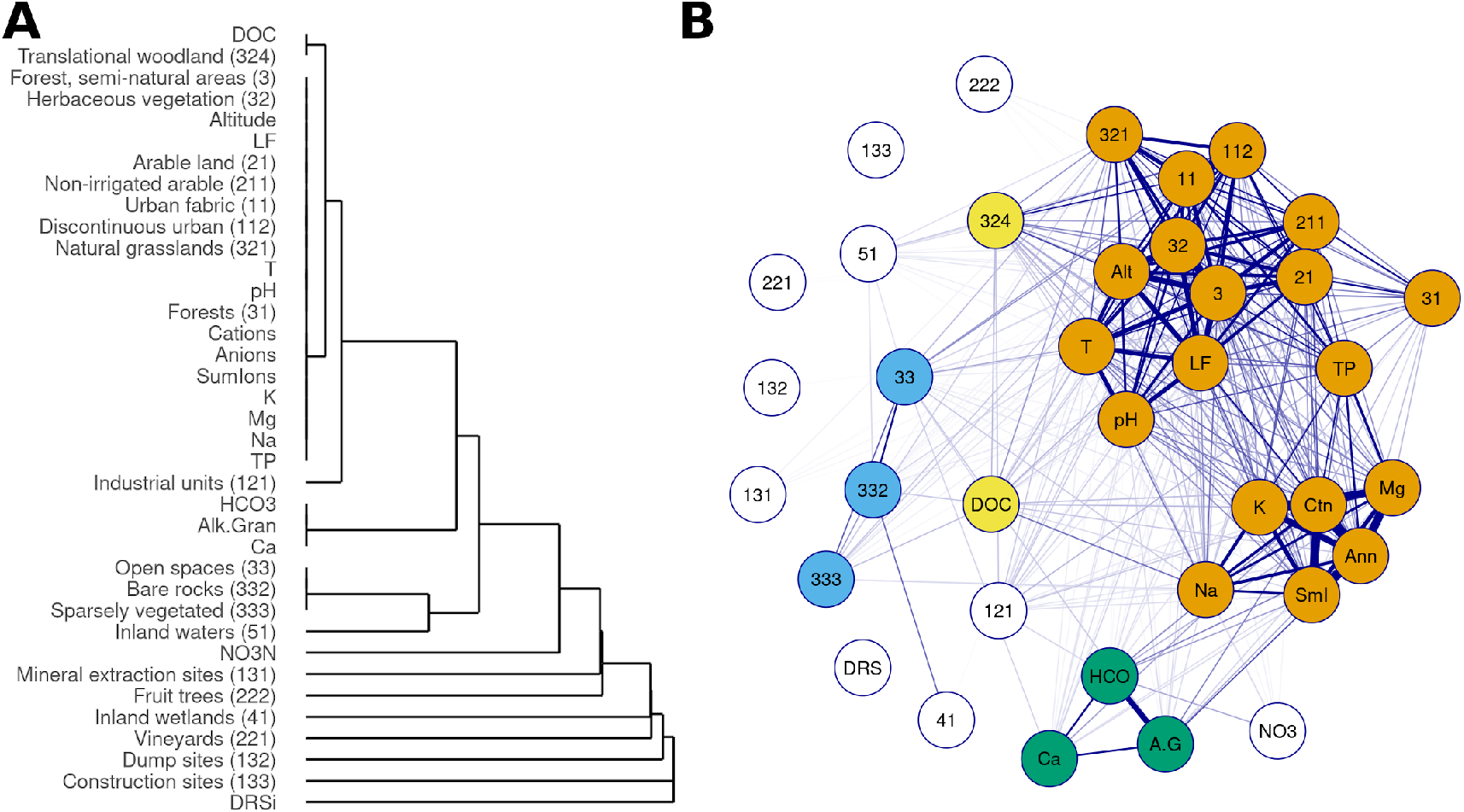
Clustering of physico-chemical and land cover parameters using the Jaccard similarity of the features’ lists of bioindicator OTUs. Visualization of the similarity matrix as **A.** dendrogram using UPGMA and **B.** force embedded graph, in which edge size represents a higher Jaccard similarity. Node coloring in **B** represents clustering of parameters in **A**. Numbers represent CLC land cover categories according to the CLC legend (see table 1). Other abbreviations: A.G/Alk.Gran – alkalinity, Alt – altitude, Ann – anions, CatSum/CtS – sum of all cation concentrations, COND/CON – conductivity, Ctn – cations, DOC – dissolved organic carbon, DRSi/DRS – dissolved reactive silica, LF – conductivity (measured in the field, SumIons/SmI – sum of all ion concentrations, TP – total phosphorus.

## 4 Discussion

The development of scalable Next-Generation Sequencing (NGS) methods has dramatically furthered the study of environmental microbiomes (Boughner and Singh, 2016, Snyder et al., 2008, Tan et al., 2015). However, technical and theoretical obstacles interfere with analyzing the wealth of data generated using NGS methods. For one, the dimensionality of microbiome datasets (i.e., the number of microbial species in ecosystems) is usually many orders of magnitude larger than the number of samples. For example, the dataset analyzed in this paper is, in terms of the number of samples, the biggest amplicon sequencing dataset for European lake microbiomes published to date, but still contains ∼1000 times more OTUs than samples. Together with the sparsity of OTU tables, this places the analyses of microbial communities at the edge of statistical feasibility, as, in such a domain, regression is ill-defined (Carr et al., 2019, Weiss et al., 2017, 2015). Furthermore, the counts of different OTUs in the OTU tables are not independent in at least two regards. Firstly, they are compositional because of technical details of the sequencing procedure (McGeoch and Chown, 1998, Quinn and Erb, 2020). Secondly, in the environment, different populations interact, through which the size of one population can, directly or indirectly, influence the size of another (Schaffer, 1981). The independence of features is, however, necessary for many statistical approaches.

In this paper, we study the relationship between lake microbiomes and the land cover surrounding the lakes while taking into account the aforementioned statistical obstacles. It is important to stress that such an analysis cannot uncover direct or functional connections without experimental validation; instead, the results presented here point to apparent relations that are present in lake ecosystems and that structure their functioning. The central analytical tool in our study is the covariation framework, in which Random Forest models are trained to approximate a projection of the microbial community composition to a one-dimensional space defined by one environmental parameter (Sperlea et al., 2021). As, in environmental, observational settings, we cannot exclude the possibility of the presence of confounding factors, we denote the relationship between the projected microbiome and the environmental parameter of interest as covariation instead of correlation to emphasize that we are not quantifying direct effects.

Our results show that the microbiome covaries to a considerable extent (*R*^2^ > 0.3) with the areas covered with forest-like vegetation (but not forest areas themselves), arable land, open spaces, plantations, and constructed environments such as urban areas and roads (see table 1, figure 2 and supplementary table 1). These results are in agreement with other recent studies of land cover and lake microbiomes (Kraemer et al., 2020, Marmen et al., 2020, Sperlea et al., 2018). On the other hand, the low covariation of the microbiome with, e.g., areas of land covered with pastures or mine and construction sites (CLC categories 23 and 13, respectively) indicate that changes in these categories of land cover are not reflected in the microbiome. On a more speculative level, these results could indicate that there are more ecological processes linking the lake microbiome to areas from the former group of land cover categories than to areas from the latter. More notable, the covariation between the microbiome and land cover categories are, generally, lower than those between the microbiome and physico-chemical parameters reported by Sperlea et al. (2021). This result is not surprising as changes in land cover do not impact lake microbiomes directly but relayed via physico-chemical parameters (Marmen et al., 2020, Saxena et al., 2015). A notable exception would be direct pollution generated through land use as, e.g., tire wear on roads and urban areas, but we do not find evidence for this in our analyses. In addition, areas that are too small to appear in the CLC dataset but that have a strong impact on lake ecology, such as a small group of trees at the shore of a lake, are not represented in our analysis.

Aggregation of similar parameters is a necessary step for the analysis of ecological processes but comes with the danger of obscuring ecologically relevant heterogeneity and, by that, losing explanatory power (Greenland and Morgenstern, 1989, King and Baker, 2010, Ulanowicz, 1986, Yodzis, 1988). For example, it has been shown that aggregation of OTUs to higher-order taxonomic groups reduces predictive power in biomonitoring settings, although this is in part due to the incompleteness of taxonomic reference databases (Chen et al., 2013, Janßen et al., 2021, Kermarrec et al., 2014, Sagova-Mareckova et al., 2021, Sperlea et al., 2021). In our results, both the aggregation of OTU tables to alpha diversity metrics (see figure 1 and supplementary figure 3) and the aggregation of land cover classes to higher-order CLC categories (see figure 2 and supplementary table 1) lead to lower *R*^2^ values. This underlines the importance of heterogeneity and non-linear processes in modelling the relationship between lake microbiomes and their habitat.

Just as the OTU counts, the environmental parameters studied here are not statistically independent but compositional and spatially autocorrelated (King et al., 2005). While, in most experimental settings, one would attempt to identify the effect of a single parameter on the study object by controlling for confounding effects via partial correlations or regression on residuals. However, spatial autocorrelation and indirect effects are significant for ecosystem functioning and integral for understanding it. They should not be considered “noise” that needs to be removed in the analysis (Freckleton, 2002, Legendre, 1993, Levins and Lewontin, 1980, Yodzis, 1988).

For example, the interdependence of ecological parameters leads to the emergence of apparent drivers of the microbial community composition, i.e., parameters that covary with the microbiome but are not necessarily directly and causally linked to it. In the context of this study, drivers appear most directly as parameters for which high numbers of microbial bioindicators have been identified. Our analysis suggests that the main driver of microbial community composition is the altitude of the lake (see table 1), which is in line with altitude being one of the determining factors of vegetation and ecosystem composition. The other parameters with high (> 500) numbers of indicators have been described as drivers of lake ecology themselves (e.g., temperature) and/or are known to be strongly correlated with altitude (as, e.g., herbaceous and forest vegetation, temperature, pollution, nutrient load), or with correlates of it (as water conductivity and pH is dependent on water temperature) (Forster et al., 2021, Urban et al., 2002). These results validate the use of microbial bioindicators for the analysis of ecosystem structure.

To gain further insights into how the microbiome responds to its environment that go beyond the concept of ecological drivers, we compared the lists of bioindicators identified for the ecological parameters studied here and in a recent publication (Sperlea et al., 2021) and visualized the results in a response map (figures 4A and B). Intuitively, environmental parameters that are close or connected with high-weighted edges in the response map elicit similar responses from the microbiome. Instead of attempting to identify direct or causal relationships, this approach allows us to analyse the structure of the ecosystem “through the eyes of the microbiome”. In principle, such an observer-based methodology allows for the integrated analysis of the effects of heterogeneous environmental parameters as long as the microbiome is affected by them.

We identify four clusters of ecological parameters in the response map (figures 4A and B), the largest of which contains the altitude, but also a wide range of land cover parameters and ion concentrations. This cluster is probably due to altitude acting as a major ecological driver. Furthermore, we find one cluster that contains the land cover parameters “Open spaces with little vegetation (33)”, “Bare Rocks (332)”, and “Sparsely vegetated areas (333)”, one cluster comprising the concentration of bicarbonate and calcium and the alkalinity, and a last one that connects the concentration of dissolved organic carbon with “Transitional woodland-shrub (324)”. The first of these clusters suggests that the sub-categorization of category 33 as present in the CLC categorization is not reflected in the lake microbiome. The second cluster might be explained by the existence of a calcium-bicarbonate equilibrium in freshwater ecosystems (Kopáček et al., 2020) and a subpopulation of the lake microbiome responding to deviations from it. The third and last cluster points towards the fact that plant matter is the principal source of dissolved organic carbon in soils (Mayer et al., 2020), and therefore, again, the functional connection between freshwater ecosystems and the landscape surrounding them.

Like environmental parameters that act as drivers of ecosystem processes, bioindicators indicative of a high number of environmental parameters, i.e., multi-target bioindicators, might be particularly sensitive with regard to environmental change. The taxonomic distribution of the multitask bioindicators for land cover categories identified here (see table 3) deserves a few words of discussion. First, the absence of Eukaryotes among the high-ranking multitask bioindicators suggests that bacterial niches can be more specific with repsect to environmental factors, making Bacteria presumably more potent and more sensitive indicators for ecosystem health. Second, the relatively low number of Alpha- and Betaproteobacteria among the multitask indicators contrasts their high abundance in a broad range of freshwater ecosystems (Šimek et al., 2013) as well as the environment-specific abundances of certain taxa among these classes (Nuy et al., 2020). Both of these findings point towards the difference between fidelity and specificity as defined in the context of the IndVal method (Cáceres and Legendre, 2009, De Cáceres et al., 2010, Dufrêne and Legendre, 1997). Third and last, the attribution of ecological functionality to these OTUs is not possible, in part because only a tiny minority of microbes have been cultured and studied to a sufficient degree (Thomas and Segata, 2019). Note that the question of how to attribute ecological function to microorganisms goes beyond the scope of this study, which is focused on the identification of microbial bioindicators.

Taken together, in this study, we analyse the relationship between land cover and lake microbiomes using machine learning methods. We propose novel methods for the analysis of relationships between ecological parameters that take the complexity of lake ecosystems into account and show that their results can be explained with regard to known ecological processes. While this study is centered around the largest lake microbiome dataset published to-date and the systematic analysis of the link between land cover and microbial ecology, it still has some limitations. For example, because the samples analyzed in this study were taken over a few days, seasonal effects are not addressed in this study. Furthermore, the fact that the underlying dataset is not longitudinal might obscure the diversity of European lakes. Therefore, further studies that adopt this methodology for large-scale microbial biomonitoring datasets are required to confirm our findings.

## Acknowledgements

Calculations on the MaRC2 high-performance computer of the University of Marburg were conducted for this research. We would like to thank René Sitt of HPC-Hessen, funded by the State Ministry of Higher Education, Research and the Arts, for the installation and maintenance of software on the MaRC2 high-performance computer. We would like to thank Marius Welzel for helping with large-scale computing. This work was supported by the LOEWE program of the State of Hesse (Germany) in the MOSLA research cluster. We also acknowledge funding by the Bauer-Foundation and Stemmler-Foundation for the project “Differential potential of metabarcoding, metatranscriptomics, and metagenomics for the assessment of lake water quality” and of the DFG project BO 3245/19-1.

## Data Accessibility

Raw sequencing data are available under the NCBI BioProject IDs PRJNA414052 and PRJNA559862.

## Author’s contributions

TS designed and performed all computational analyses, JPS provided the OSM dataset, HD helped with the network analyses, DB performed the amplicon sequence analysis with Natrix, DB and JB provided the sequencing datasets, JB, GH, and DH supervised the study. All authors discussed the results and wrote and revised the manuscript.

## References

C. R. Allen, D. G. Angeler, A. S. Garmestani, L. H. Gunderson, and C. S. Holling. Panarchy: Theory and application. Ecosystems, 17(4):578–589, 1 2014. doi: 10.1007/s10021-013-9744-2. URLhttp://dx.doi.org/10.1007/s10021-013-9744-2.

S. F. Altschul, W. Gish, W. Miller, E. W. Myers, and D. J. Lipman. Basic local alignment search tool. Journal of Molecular Biology, 215(3):403–410, oct 1990. doi: 10.1016/s0022-2836(05)80360-2. URL https://doi.org/10.1016/s0022-2836(05)80360-2.

S. Andrews. FASTQC. A quality control tool for high throughput sequence data, 2010.

D. Berry and S. Widder. Deciphering microbial interactions and detecting keystone species with co-occurrence networks. Frontiers in Microbiology, 5, may 2014. doi: 10.3389/fmicb.2014.00219. URL https://doi.org/10.3389/fmicb.2014.00219.

S. Birk, W. Bonne, A. Borja, S. Brucet, A. Courrat, S. Poikane, A. Solimini, W. van de Bund, N. Zampoukas, and D. Hering. Three hundred ways to assess Europe’s surface waters: An almost complete overview of biological methods to implement the Water Framework Directive. Ecological Indicators, 18:31–41, jul 2012. doi: 10.1016/j.ecolind.2011.10.009. URL https://doi.org/10.1016/j.ecolind.2011.10.009.

C. Bock, M. Jensen, D. Forster, S. Marks, J. Nuy, R. Psenner, D. Beisser, and J. Boenigk. Factors shaping community patterns of protists and bacteria on a european scale. Environmental Microbiology, 22(6):2243–2260, mar 2020. doi: 10.1111/1462-2920.14992. URL https://doi.org/10.1111/1462-2920.14992.

J. Boenigk, S. Wodniok, C. Bock, D. Beisser, C. Hempel, L. Grossmann, A. Lange, and M. Jensen. Geographic distance and mountain ranges structure freshwater protist communities on a european scale. Metabarcoding and Metagenomics, 2:e21519, jan 2018. doi: 10.3897/mbmg.2.21519. URL https://doi.org/10.3897/mbmg.2.21519.

L. A. Boughner and P. Singh. Microbial ecology: Where are we now? Postdoc Journal, 4(11), nov 2016. doi: 10.14304/surya.jpr.v4n11.2. URL https://doi.org/10.14304/surya.jpr.v4n11.2.

J. R. Bray and J. T. Curtis. An ordination of the upland forest communities of southern wisconsin. Ecological Monographs, 27(4):325–349, oct 1957. doi: 10.2307/1942268. URL https://doi.org/10.2307/1942268.

L. Breiman. Random forests. Machine Learning, 45(1):5–32, 2001. doi: 10.1023/a:1010933404324. URL https://doi.org/10.1023/a:1010933404324.

A. Carr, C. Diener, N. S. Baliga, and S. M. Gibbons. Use and abuse of correlation analyses in microbial ecology. The ISME Journal, 13(11):2647–2655, jun 2019. doi: 10.1038/s41396-019-0459-z. URL https://doi.org/10.1038/s41396-019-0459-z.

A. Chao and L. Jost. Estimating diversity and entropy profiles via discovery rates of new species. Methods in Ecology and Evolution, 6(8):873–882, feb 2015. doi: 10.1111/2041-210x.12349. URL https://doi.org/10.1111/2041-210x.12349.

W. Chen, C. K. Zhang, Y. Cheng, S. Zhang, and H. Zhao. A comparison of methods for clustering 16S rrna sequences into otus. PLoS ONE, 8(8):e70837, aug 2013. doi: 10.1371/journal.pone.0070837. URL https://doi.org/10.1371/journal.pone.0070837.

R. R. Colwell. Microbial diversity: the importance of exploration and conservation. Journal of Industrial Microbiology and Biotechnology, 18(5):302–307, 5 1997. doi: 10.1038/sj.jim.2900390. URL http://dx.doi.org/10.1038/sj.jim.2900390.

E. F. Connor and D. Simberloff. The assembly of species communities: Chance or competition? Ecology, 60 (6):1132–1140, 1979. ISSN 00129658, 19399170. URL http://www.jstor.org/stable/1936961.

T. Cordier, D. Forster, Y. Dufresne, C. I. M. Martins, T. Stoeck, and J. Pawlowski. Supervised machine learning outperforms taxonomy-based environmental DNA metabarcoding applied to biomonitoring. Molecular Ecology Resources, 18(6):1381–1391, aug 2018. doi: 10.1111/1755-0998.12926. URL https://doi.org/10.1111/1755-0998.12926.

G. Csardi and T. Nepusz. The igraph software package for complex network research. InterJournal, Complex Systems:1695, 2006. URL https://igraph.org.

M. D. Cáceres and P. Legendre. Associations between species and groups of sites: indices and statistical inference. Ecology, 90(12):3566–3574, 2009. doi:https://doi.org/10.1890/08-1823.1.

A. Daly, J. Baetens, and B. D. Baets. Ecological diversity: Measuring the unmeasurable. Mathematics, 6(7): 119, jul 2018. doi: 10.3390/math6070119. URL https://doi.org/10.3390/math6070119.

M. De Cáceres, P. Legendre, and M. Moretti. Improving indicator species analysis by combining groups of sites. Oikos, 119(10):1674–1684, 9 2010. doi: 10.1111/j.1600-0706.2010.18334.x. URL http://dx.doi.org/10.1111/j.1600-0706.2010.18334.x.

A. de Vries and B. D. Ripley. ggdendro: Create Dendrograms and Tree Diagrams Using ‘ggplot2’, 2020. URL https://CRAN.R-project.org/package=ggdendro. R package version 0.1.22.

T. Dobzhansky. Are naturalists old-fashioned? The American Naturalist, 100(915):541–550, 9 1966. doi: 10.1086/282448. URL http://dx.doi.org/10.1086/282448.

K. M. Docherty and J. L. M. Gutknecht. The role of environmental microorganisms in ecosystem responses to global change: current state of research and future outlooks. Biogeochemistry, 109(1-3):1–6, sep 2011. doi: 10.1007/s10533-011-9614-y. URL https://doi.org/10.1007/s10533-011-9614-y.

M. Dufrêne and P. Legendre. Species assemblages and indicator species: The need for a flexible asymmetrical approach. Ecological Monographs, 67(3):345–366, aug 1997. doi: 10.1890/0012-9615(1997)067[0345:saaist]2.0.co;2.

R. C. Edgar, B. J. Haas, J. C. Clemente, C. Quince, and R. Knight. UCHIME improves sensitivity and speed of chimera detection. Bioinformatics, 27(16):2194–2200, jun 2011. doi: 10.1093/bioinformatics/btr381. URL https://doi.org/10.1093/bioinformatics/btr381.

S. Epskamp, A. O. J. Cramer, L. J. Waldorp, V. D. Schmittmann, and D. Borsboom. qgraph: Network visualizations of relationships in psychometric data. Journal of Statistical Software, 48(4):1–18, 2012.

European Union. Copernicus land monitoring service 2012. European Environment Agency (EEA), 2012.

D. Forster, Z. Qu, G. Pitsch, E. P. Bruni, B. Kammerlander, T. Pröschold, B. Sonntag, T. Posch, and T. Stoeck. Lake ecosystem robustness and resilience inferred from a climate-stressed protistan plankton network. Microorganisms, 9(3):549, 3 2021. doi: 10.3390/microorganisms9030549. URL http://dx.doi.org/10.3390/microorganisms9030549.

R. P. Freckleton. On the misuse of residuals in ecology: regression of residuals vs. multiple regression. Journal of Animal Ecology, 71(3):542–545, 5 2002. doi: 10.1046/j.1365-2656.2002.00618.x. URL http://dx.doi.org/10.1046/j.1365-2656.2002.00618.x.

J. Friedman and E. J. Alm. Inferring correlation networks from genomic survey data. PLoS Computational Biology, 8(9):e1002687, sep 2012. doi: 10.1371/journal.pcbi.1002687. URL https://doi.org/10.1371/journal.pcbi.1002687.

R. C. Gatti, B. Fath, W. Hordijk, S. Kauffman, and R. Ulanowicz. Niche emergence as an autocatalytic process in the evolution of ecosystems. Journal of Theoretical Biology, 454:110–117, 2018. doi: https://doi.org/10.1016/j.jtbi.2018.05.038. URL http://www.sciencedirect.com/science/article/pii/S0022519318302856.

D. G. Green and S. Sadedin. Interactions matter—complexity in landscapes and ecosystems. Ecological Complexity, 2(2):117–130, jun 2005. doi: 10.1016/j.ecocom.2004.11.006. URL https://doi.org/10.1016/j.ecocom.2004.11.006.

S. Greenland and H. Morgenstern. Ecological bias, confounding, and effect modification. International Journal of Epidemiology, 18(1):269–274, 1989. doi: 10.1093/ije/18.1.269. URL http://dx.doi.org/10.1093/ije/18.1.269.

M. Han, M. Dsouza, C. Zhou, H. Li, J. Zhang, C. Chen, Q. Yao, C. Zhong, H. Zhou, J. A. Gilbert, Z. Wang, and K. Ning. Agricultural risk factors influence microbial ecology in honghu lake. GenomicsProteomics & Bioinformatics, 17(1):76–90, feb 2019. doi: 10.1016/j.gpb.2018.04.008. URL https://doi.org/10.1016/j.gpb.2018.04.008.

G. Hattab, T.-M. Rhyne, and D. Heider. Ten simple rules to colorize biological data visualization. PLOS Computational Biology, 16(10):e1008259, 10 2020. doi: 10.1371/journal.pcbi.1008259. URL http://dx.doi.org/10.1371/journal.pcbi.1008259.

D. Hering, A. Borja, J. Carstensen, L. Carvalho, M. Elliott, C. K. Feld, A.-S. Heiskanen, R. K. Johnson, J. Moe, and D. Pont. The European Water Framework Directive at the age of 10: A critical review of the achievements with recommendations for the future. Science of The Total Environment, 408(19):4007–4019, Sep 2010. doi: 10.1016/j.scitotenv.2010.05.031. URL https://doi.org/10.1016/j.scitotenv.2010.05.031.

Intergovernmental Platform on Biodiversity and Ecosystem Services (IPBES). Media release: Nature’s dangerous decline ‘unprecedented’species extinction rates ‘accelerating’, 2019. URL https://www.ipbes.net/sites/default/files/downloads/20180322_ipbes6_media_release_regional_assessments_en.pdf.

R. Janßen, A. J. Beck, J. Werner, O. Dellwig, J. Alneberg, B. Kreikemeyer, E. Maser, C. Böttcher, E. P. Achterberg, A. F. Andersson, and M. Labrenz. Machine learning predicts the presence of 2,4,6-trinitrotoluene in sediments of a baltic sea munitions dumpsite using microbial community compositions. Frontiers in Microbiology, 12, 9 2021. doi: 10.3389/fmicb.2021.626048. URL http://dx.doi.org/10.3389/fmicb.2021.626048.

J. Karlsson, A. Jonsson, and M. Jansson. Productivity of high-latitude lakes: climate effect inferred from altitude gradient. Global Change Biology, 11(5):710–715, may 2005. doi: 10.1111/j.1365-2486.2005.00945.x. URL https://doi.org/10.1111/j.1365-2486.2005.00945.x.

L. Kermarrec, A. Franc, F. Rimet, P. Chaumeil, J.-M. Frigerio, J.-F. Humbert, and A. Bouchez. A next-generation sequencing approach to river biomonitoring using benthic diatoms. Freshwater Science, 33(1): 349–363, mar 2014. doi: 10.1086/675079. URL https://doi.org/10.1086/675079.

R. S. King and M. E. Baker. Considerations for analyzing ecological community thresholds in response to anthropogenic environmental gradients. Journal of the North American Benthological Society, 29(3): 998–1008, 9 2010. doi: 10.1899/09-144.1. URL http://dx.doi.org/10.1899/09-144.1.

R. S. King, M. E. Baker, D. F. Whigham, D. E. Weller, T. E. Jordan, P. F. Kazyak, and M. K. Hurd. Spatial considerations for linking watershed land cover to ecological indicators in streams. Ecological Applications, 15, 2005. doi: https://doi.org/10.1890/04-0481. URL https://esajournals.onlinelibrary.wiley.com/doi/abs/10.1890/04-0481.

J. Kopáček, J. Hejzlar, F. Oulehle, P. Porcal, G. A. Weyhenmeyer, and S. A. Norton. Disruptions and re-establishment of the calcium-bicarbonate equilibrium in freshwaters. Science of The Total Environment, 743:140626, 11 2020. doi: 10.1016/j.scitotenv.2020.140626. URL http://dx.doi.org/10.1016/j.scitotenv.2020.140626.

S. A. Kraemer, N. B. da Costa, B. J. Shapiro, M. Fradette, Y. Huot, and D. A. Walsh. A large-scale assessment of lakes reveals a pervasive signal of land use on bacterial communities. The ISME Journal, 14:3011–3023, aug 2020. doi: 10.1038/s41396-020-0733-0. URL https://doi.org/10.1038/s41396-020-0733-0.

M. Kuhn. Building predictive models in R using the caret package. Journal of Statistical Software, 28(5), 2008. doi: 10.18637/jss.v028.i05. URL https://doi.org/10.18637/jss.v028.i05.

S. Kullback and R. A. Leibler. On information and sufficiency. The Annals of Mathematical Statistics, 22(1): 79–86, mar 1951. doi: 10.1214/aoms/1177729694. URL https://doi.org/10.1214/aoms/1177729694.

Z. D. Kurtz, C. L. Müller, E. R. Miraldi, D. R. Littman, M. J. Blaser, and R. A. Bonneau. Sparse and compositionally robust inference of microbial ecological networks. PLOS Computational Biology, 11(5): e1004226, may 2015. doi: 10.1371/journal.pcbi.1004226. URL https://doi.org/10.1371/journal.pcbi.1004226.

P. B. Landres, J. Verner, and J. W. Thomas. Ecological uses of vertebrate indicator species: A critique. Conservation Biology, 2(4):316–328, 12 1988. doi: 10.1111/j.1523-1739.1988.tb00195.x. URL http://dx.doi.org/10.1111/j.1523-1739.1988.tb00195.x.

A. Lange, S. Jost, D. Heider, C. Bock, B. Budeus, E. Schilling, A. Strittmatter, J. Boenigk, and D. Hoffmann. Ampliconduo: A split-sample filtering protocol for high-throughput amplicon sequencing of microbial communities. PLOS ONE, 10(11):e0141590, nov 2015. doi: 10.1371/journal.pone.0141590. URL https://doi.org/10.1371/journal.pone.0141590.

P. Legendre. Spatial autocorrelation: Trouble or new paradigm? Ecology, 74(6):1659–1673, 9 1993. doi: 10.2307/1939924. URL http://dx.doi.org/10.2307/1939924.

P. Legendre and E. D. Gallagher. Ecologically meaningful transformations for ordination of species data. Oecologia, 129(2):271–280, oct 2001. doi: 10.1007/s004420100716. URL https://doi.org/10.1007/s004420100716.

S. A. Levin. Ecosystems and the biosphere as complex adaptive systems. Ecosystems, 1(5):431–436, sep 1998. doi: 10.1007/s100219900037. URL https://doi.org/10.1007/s100219900037.

R. Levins and R. Lewontin. Dialectics and reductionism in ecology. Synthese, 43(1):47–78, jan 1980. doi: 10.1007/bf00413856. URL https://doi.org/10.1007/bf00413856.

R. Logares, S. V. Tesson, B. Canbäck, M. Pontarp, K. Hedlund, and K. Rengefors. Contrasting prevalence of selection and drift in the community structuring of bacteria and microbial eukaryotes. Environmental Microbiology, may 2018. doi: 10.1111/1462-2920.14265. URL https://doi.org/10.1111/1462-2920.14265.

F. Mahé, T. Rognes, C. Quince, C. de Vargas, and M. Dunthorn. Swarm: robust and fast clustering method for amplicon-based studies. PeerJ, 2:e593, sep 2014. doi: 10.7717/peerj.593. URL https://doi.org/10.7717/peerj.593.

S. Marmen, L. Blank, A. Al-Ashhab, A. Malik, L. Ganzert, M. Lalzar, H.-P. Grossart, and D. Sher. The role of land use types and water chemical properties in structuring the microbiomes of a connected lake system. Frontiers in Microbiology, 11, feb 2020. doi: 10.3389/fmicb.2020.00089. URL https://doi.org/10.3389/fmicb.2020.00089.

G. Martin, C. Dang, E. Morrissey, J. Hubbart, E. Kellner, C. Kelly, K. Stephan, and Z. Freedman. Stream sediment bacterial communities exhibit temporally-consistent and distinct thresholds to land use change in a mixed-use watershed. FEMS Microbiology Ecology, 97(2), 12 2020. doi: 10.1093/femsec/fiaa256. URL http://dx.doi.org/10.1093/femsec/fiaa256.

A. P. Masella, A. K. Bartram, J. M. Truszkowski, D. G. Brown, and J. D. Neufeld. Pandaseq: paired-end assembler for illumina sequences. BMC Bioinformatics, 13(1):31, 2012. doi: 10.1186/1471-2105-13-31. URL https://doi.org/10.1186/1471-2105-13-31.

M. Mayer, C. E. Prescott, W. E. Abaker, L. Augusto, L. Cécillon, G. W. Ferreira, J. James, R. Jandl, K. Katzensteiner, J.-P. Laclau, J. Laganière, Y. Nouvellon, D. Paré, J. A. Stanturf, E. I. Vanguelova, and L. Vesterdal. Tamm review: Influence of forest management activities on soil organic carbon stocks:A knowledge synthesis. Forest Ecology and Management, 466:118127, 2020. ISSN 0378-1127. doi: https://doi.org/10.1016/j.foreco.2020.118127. URL https://www.sciencedirect.com/science/article/pii/S0378112720300268.

M. A. McGeoch and S. L. Chown. Scaling up the value of bioindicators. Trends in Ecology & Evolution, 13(2): 46–47, 2 1998. doi: 10.1016/s0169-5347(97)01279-2. URL http://dx.doi.org/10.1016/S0169-5347(97)01279-2.

J. K. Nuy, M. Hoetzinger, M. W. Hahn, D. Beisser, and J. Boenigk. Ecological differentiation in two major freshwater bacterial taxa along environmental gradients. Frontiers in Microbiology, 11, feb 2020. doi: 10.3389/fmicb.2020.00154. URL https://doi.org/10.3389/fmicb.2020.00154.

E.P. Odum. Ecology - A Bridge Between Science And Society. Sinauer Associates Incorporated, 1997. ISBN 9780878936304.

J. Oksanen, F. G. Blanchet, M. Friendly, R. Kindt, P. Legendre, D. McGlinn, P. R. Minchin, R. B. O’Hara, G. L. Simpson, P. Solymos, M. H. H. Stevens, E. Szoecs, and H. Wagner. vegan: Community Ecology Package, 2019. URL https://CRAN.R-project.org/package=vegan. R package version 2.5-6.

R. V. O’Neill, C. T. Hunsaker, K. B. Jones, K. H. Riitters, J. D. Wickham, P. M. Schwartz, I. A. Goodman, B. L. Jackson, and W. S. Baillargeon. Monitoring environmental quality at the landscape scale. BioScience, 47(8):513–519, 9 1997. doi: 10.2307/1313119. URL http://dx.doi.org/10.2307/1313119.

OpenStreetMap contributors. Planet dump retrieved from https://planet.osm.org. https://www.openstreetmap.org, 2017.

QGIS Development Team. QGIS Geographic Information System. QGIS Association, 2020. URL https://www.qgis.org.

C. Quast, E. Pruesse, P. Yilmaz, J. Gerken, T. Schweer, P. Yarza, J. Peplies, and F. O. Glöckner. The SILVA ribosomal RNA gene database project: improved data processing and web-based tools. Nucleic Acids Research, 41(D1):D590–D596, Nov. 2012. doi: 10.1093/nar/gks1219. URL https://doi.org/10.1093/nar/gks1219.

T. P. Quinn and I. Erb. Amalgams: data-driven amalgamation for the dimensionality reduction of compositional data. NAR Genomics and Bioinformatics, 2(4), 10 2020. doi: 10.1093/nargab/lqaa076. URL http://dx.doi.org/10.1093/nargab/lqaa076.

M. Sagova-Mareckova, J. Boenigk, A. Bouchez, K. Cermakova, T. Chonova, T. Cordier, U. Eisendle, T. Elersek, S. Fazi, T. Fleituch, L. Frühe, M. Gajdosova, N. Graupner, A. Haegerbaeumer, A.-M. Kelly, J. Kopecky, F. Leese, P. N. oges, S. Orlic, K. Panksep, J. Pawlowski, A. Petrusek, J. Piggott, J. Rusch, R. Salis, J. Schenk, K. Simek, A. Stovicek, D. Strand, M. Vasquez, T. Vrålstad, S. Zlatkovic, M. Zupancic, and T. Stoeck. Expanding ecological assessment by integrating microorganisms into routine fresh-water biomonitoring. Water Research, 191:116767, 3 2021. doi: 10.1016/j.watres.2020.116767. URL http://dx.doi.org/10.1016/j.watres.2020.116767.

G. Saxena, E. M. Marzinelli, N. N. Naing, Z. He, Y. Liang, L. Tom, S. Mitra, H. Ping, U. M. Joshi, S. Reuben, K. C. Mynampati, S. Mishra, S. Umashankar, J. Zhou, G. L. Andersen, S. Kjelleberg, and S. Swarup. Ecogenomics reveals metals and land-use pressures on microbial communities in the waterways of a megacity. Environmental Science & Technology, 49(3):1462–1471, jan 2015. doi: 10.1021/es504531s. URL https://doi.org/10.1021/es504531s.

W. M. Schaffer. Ecological abstraction: The consequences of reduced dimensionality in ecological models. Ecological Monographs, 51(4):383–401, 12 1981. doi: 10.2307/2937321. URL http://dx.doi.org/10.2307/2937321.

K. Schliep. phangorn: phylogenetic analysis in r. Bioinformatics, 27(4):592–593, 2011. URL https://doi.org/10.1093/bioinformatics/btq706.

R. Schmieder and R. Edwards. Quality control and preprocessing of metagenomic datasets. Bioinformatics, 27(6):863–864, jan 2011. doi: 10.1093/bioinformatics/btr026. URL https://doi.org/10.1093/bioinformatics/btr026.

D. Simberloff. Flagships, umbrellas, and keystones: Is single-species management passé in the landscape era? Biological Conservation, 83(3):247–257, 3 1998. doi: 10.1016/s0006-3207(97)00081-5. URL http://dx.doi.org/10.1016/S0006-3207(97)00081-5.

L. A. S. Snyder, N. Loman, M. J. Pallen, and C. W. Penn. Next-generation sequencing—the promise and perils of charting the great microbial unknown. Microbial Ecology, 57(1):1–3, nov 2008. doi: 10.1007/s00248-008-9465-9. URL https://doi.org/10.1007/s00248-008-9465-9.

X.-P. Song, M. C. Hansen, S. V. Stehman, P. V. Potapov, A. Tyukavina, E. F. Vermote, and J. R. Townshend. Global land change from 1982 to 2016. Nature, 560(7720):639–643, aug 2018. doi: 10.1038/s41586-018-0411-9. URL https://doi.org/10.1038/s41586-018-0411-9.

T. Sperlea, S. Füser, J. Boenigk, and D. Heider. SEDE-GPS: socio-economic data enrichment based on GPS information. BMC Bioinformatics, 19(440), nov 2018. doi: 10.1186/s12859-018-2419-4. URL https://doi.org/10.1186/s12859-018-2419-4.

T. Sperlea, N. Kreuder, D. Beisser, G. Hattab, J. Boenigk, and D. Heider. Quantification of the covariation of lake microbiomes and environmental variables using a machine learning-based framework. Molecular Ecology, 30(9):2131–2144, 5 2021. doi: 10.1111/mec.15872. URL http://dx.doi.org/10.1111/mec.15872.

M. J. Steinbauer, J.-A. Grytnes, G. Jurasinski, A. Kulonen, J. Lenoir, H. Pauli, C. Rixen, M. Winkler, M. Bardy-Durchhalter, E. Barni, A. D. Bjorkman, F. T. Breiner, S. Burg, P. Czortek, M. A. Dawes, A. Delimat, S. Dullinger, B. Erschbamer, V. A. Felde, O. Fernández-Arberas, K. F. Fossheim, D. Gómez-García, D. Georges, E. T. Grindrud, S. Haider, S. V. Haugum, H. Henriksen, M. J. Herreros, B. Jaroszewicz, F. Jaroszynska, R. Kanka, J. Kapfer, K. Klanderud, I. Kühn, A. Lamprecht, M. Matteodo, U. M. di Cella, S. Normand, A. Odland, S. L. Olsen, S. Palacio, M. Petey, V. Piscová, B. Sedlakova, K. Steinbauer, V. StĂśckli, J.-C. Svenning, G. Teppa, J.-P. Theurillat, P. Vittoz, S. J. Woodin, N. E. Zimmermann, and S. Wipf. Accelerated increase in plant species richness on mountain summits is linked to warming. Nature, 556(7700):231–234, apr 2018. doi: 10.1038/s41586-018-0005-6. URL https://doi.org/10.1038/s41586-018-0005-6.

L. Stone and A. Roberts. The checkerboard score and species distributions. Oecologia, 85(1):74–79, nov 1990. doi: 10.1007/bf00317345. URL https://doi.org/10.1007/bf00317345.

B. Tan, C. Ng, J. P. Nshimyimana, L. L. Loh, K. Y.-H. Gin, and J. R. Thompson. Next-generation sequencing (NGS) for assessment of microbial water quality: current progress, challenges and future opportunities. Frontiers in Microbiology, 6, sep 2015. doi: 10.3389/fmicb.2015.01027. URL https://doi.org/10.3389/fmicb.2015.01027.

A. M. Thomas and N. Segata. Multiple levels of the unknown in microbiome research. BMC Biology, 17, jun 2019. doi: 10.1186/s12915-019-0667-z.

R. E. Ulanowicz. Growth and Development. Springer New York, 1986. doi: 10.1007/978-1-4612-4916-0. URL https://doi.org/10.1007/978-1-4612-4916-0.

D. Urban, S. Goslee, K. Pierce, and T. Lookingbill. Extending community ecology to landscapes. Écoscience, 9(2):200–212, 1 2002. doi: 10.1080/11956860.2002.11682706. URL http://dx.doi.org/10.1080/11956860.2002.11682706.

K. Šimek, V. Kasalický, J. Jezbera, K. Horňák, J. Nedoma, M. W. Hahn, D. Bass, S. Jost, and J. Boenigk. Differential freshwater flagellate community response to bacterial food quality with a focus on limno-habitans bacteria. The ISME Journal, 7(8):1519–1530, apr 2013. doi: 10.1038/ismej.2013.57. URL https://doi.org/10.1038/ismej.2013.57.

B. Wang, X. Zheng, H. Zhang, F. Xiao, Z. He, and Q. Yan. Keystone taxa of water microbiome respond to environmental quality and predict water contamination. Environmental Research, 187:109666, aug 2020. doi: 10.1016/j.envres.2020.109666. URL https://doi.org/10.1016/j.envres.2020.109666.

N. S. Webster, M. Wagner, and A. P. Negri. Microbial conservation in the Anthropocene. Environmental Microbiology, 20(6):1925–1928, may 2018. doi: 10.1111/1462-2920.14124. URL https://doi.org/10.1111/1462-2920.14124.

S. Weiss, Z. Z. Xu, S. Peddada, A. Amir, K. Bittinger, A. Gonzalez, C. Lozupone, J. R. Zaneveld, Y. Vázquez-Baeza, A. Birmingham, E. R. Hyde, and R. Knight. Normalization and microbial differential abundance strategies depend upon data characteristics. Microbiome, 5(1), mar 2017. doi: 10.1186/s40168-017-0237-y. URL https://doi.org/10.1186/s40168-017-0237-y.

S. J. Weiss, Z. Xu, A. Amir, S. Peddada, K. Bittinger, A. Gonzalez, C. Lozupone, J. R. Zaneveld, Y. Vazquez-Baeza, A. Birmingham, and R. Knight. Effects of library size variance sparsity and compositionality on the analysis of microbiome data. PeerJ PrePrints, 2015. doi: 10.7287/peerj.preprints.1157v1.

M. Welzel, A. Lange, D. Heider, M. Schwarz, B. Freisleben, M. Jensen, J. Boenigk, and D. Beisser. Natrix: a Snakemake-based workflow for processing, clustering, and taxonomically assigning amplicon sequencing reads. BMC Bioinformatics, 21(526), nov 2020. doi: 10.1186/s12859-020-03852-4. URL https://doi.org/10.1186/s12859-020-03852-4.

H. Wickham. ggplot2: Elegant Graphics for Data Analysis. Springer-Verlag New York, 2016. URL https://ggplot2.tidyverse.org.

C. E. Williamson, W. Dodds, T. K. Kratz, and M. A. Palmer. Lakes and streams as sentinels of environmental change in terrestrial and atmospheric processes. Frontiers in Ecology and the Environment, 6(5):247–254, jun 2008. doi: 10.1890/070140. URL https://doi.org/10.1890/070140.

P. Yodzis. The indeterminacy of ecological interactions as perceived through perturbation experiments. Ecology, 69(2):508–515, 4 1988. doi: 10.2307/1940449. URL http://dx.doi.org/10.2307/1940449.

