## Supplementary material for "The relationship between land cover and microbial community composition in European lakes": si.pdf

### Supplementary tables

**Supplementary table 1**: `st_1_covar_clc.csv`
Results for the covariation framework ($R^2$, upper and lower bounds of the confidence intervals) for all categories at all levels of the CORINE Land Cover hierarchy at radii between 1 and 10 km.
**Supplementary table 2**: `st_2_bioindicators_w_taxonomy.csv`
A list of all bioindicators identified for land cover classes in this study. The table is sorted by the amount of times an OTU has been identified for land cover classes (column *Freq*), and contains information on the number of times the same OTU has been identified for physico-chemical parameters (column *Freq_physchem*), which parameters those are (column *targets*) as well as the taxonomic annotation for the OTUs.
**Supplementary table 3**: `st_3_bioindicators_network.sif`
The bioindicator network generated in this study in the sif format. Note that as edges are not weighted in this graph, the third column only contains values of 1; edges that are not present in the network are also omitted in the file.

### Supplementary figures

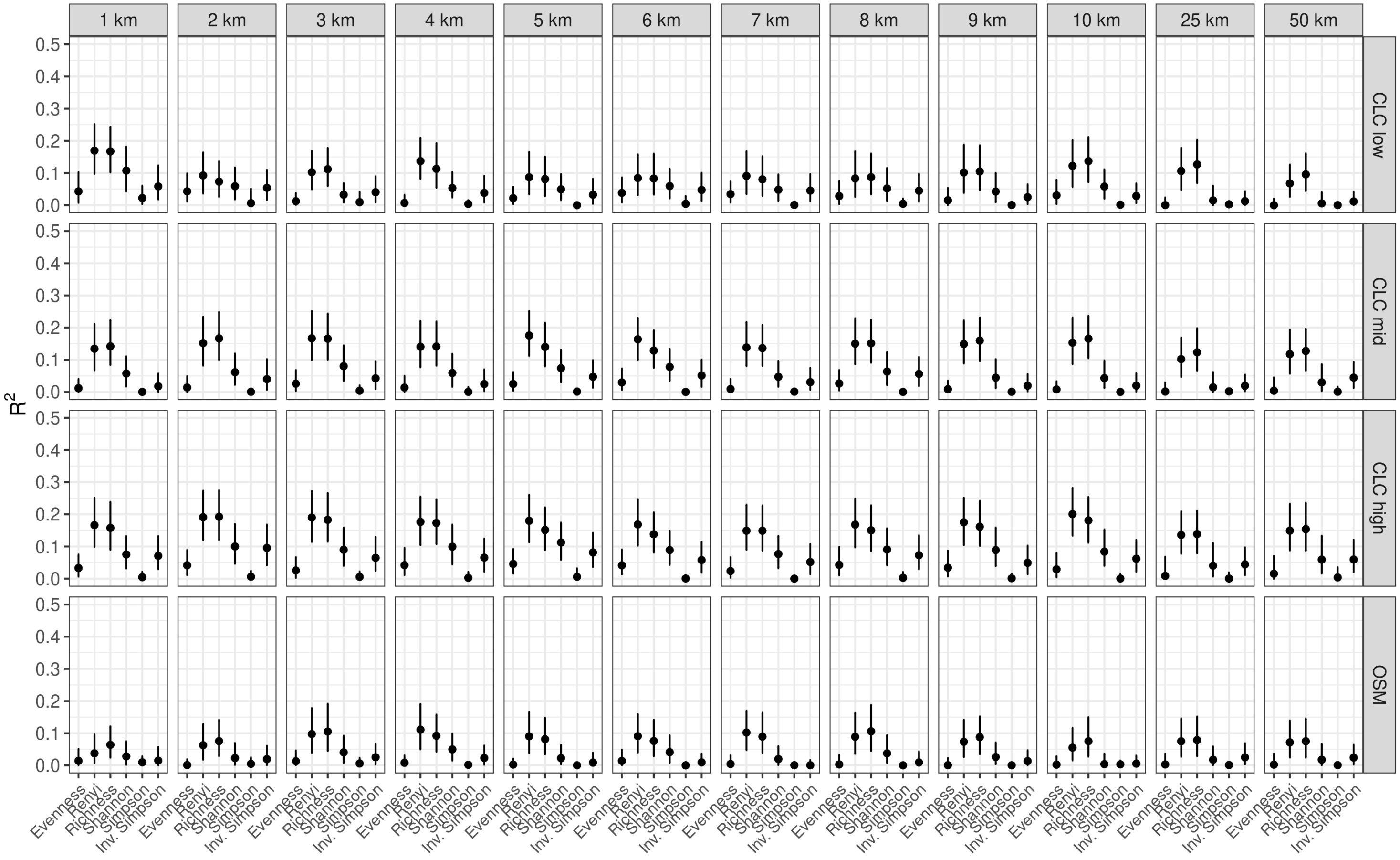


Supplementary Figure 1: Evaluation results of RF models trained to predict biodiversity metrics from land cover for all combinations of land cover dataset, radius and biodiversity metric. Renyi entropy is given for $\alpha = 0.5$; for the results at other values of $\alpha$, see supplementary figure 2.

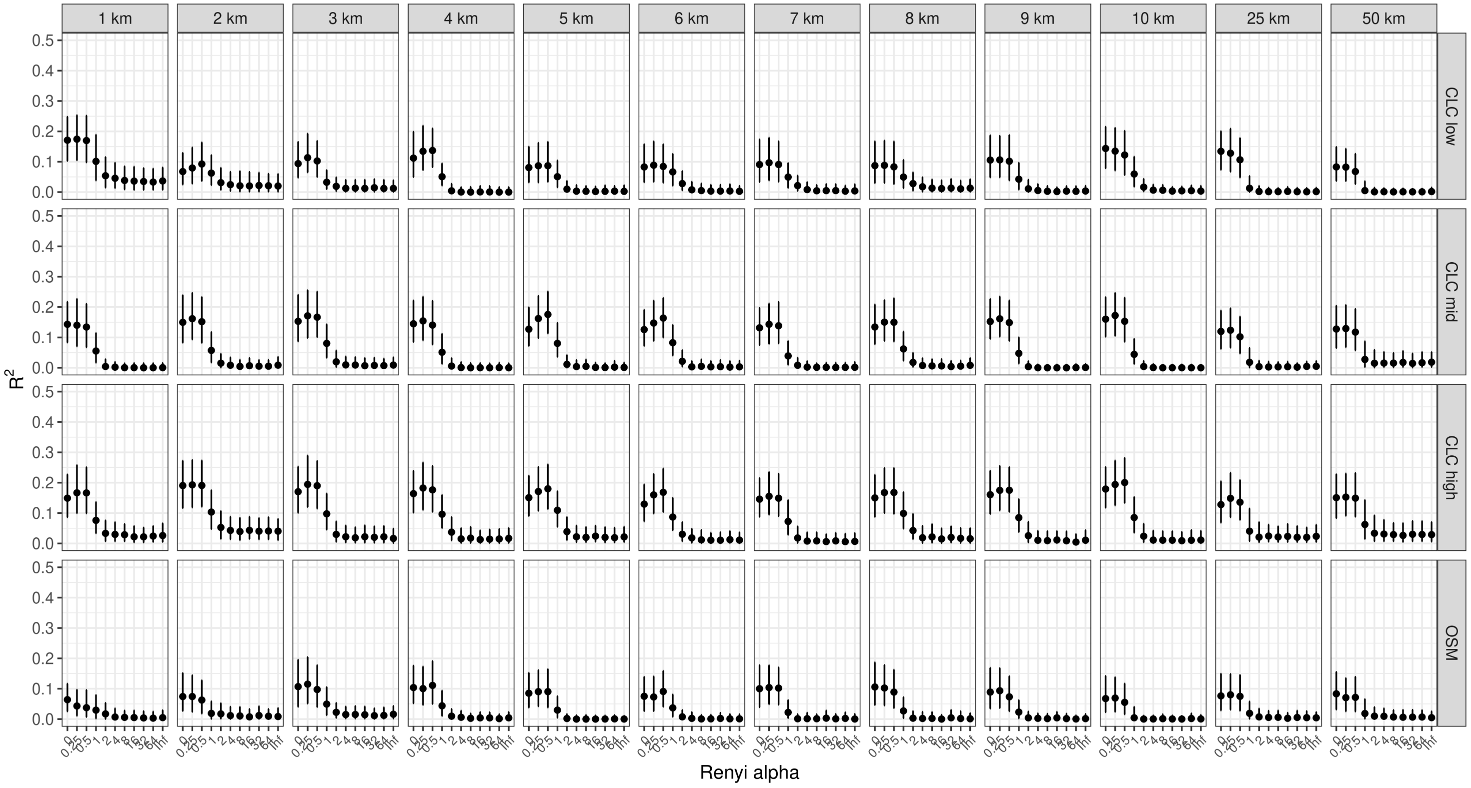


Supplementary Figure 2: Evaluation results of RF models trained to predict Renyi diversity from land cover for all combinations of land cover dataset, radius and value of $\alpha$.

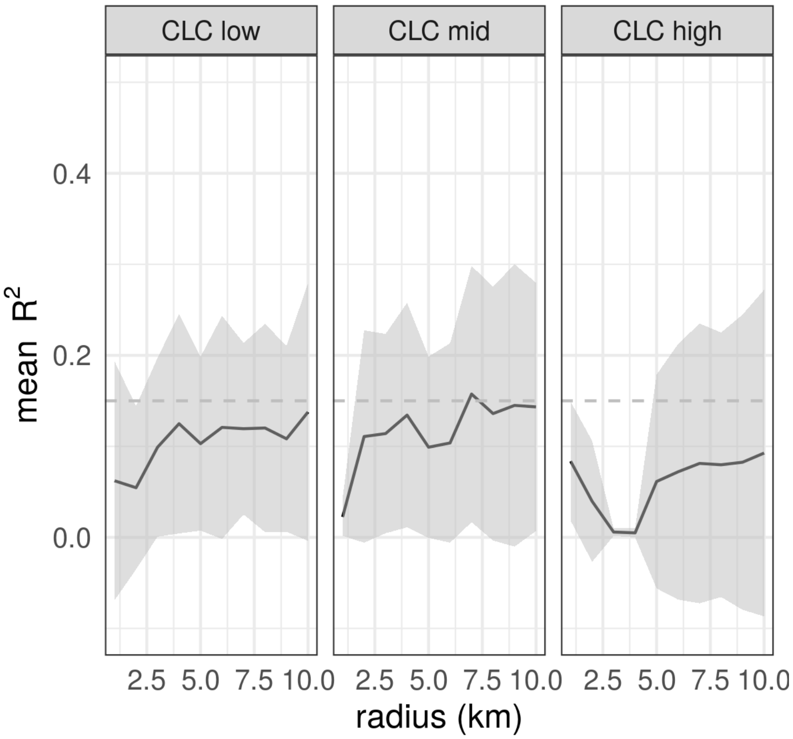


Supplementary Figure 3: Average covariation of all land cover categories at the three levels of CLC category hierarchy. Shaded area represents standard deviation.

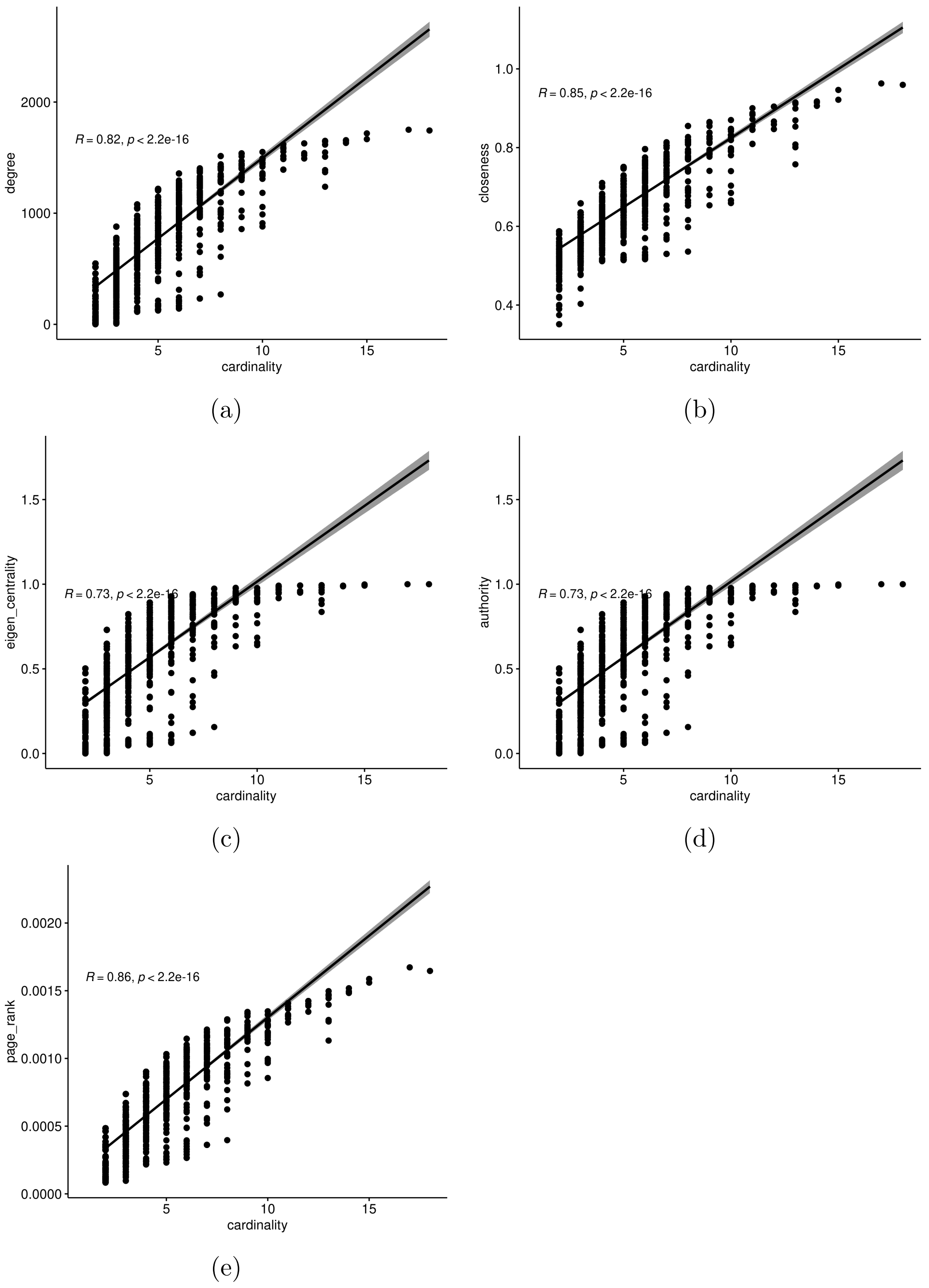


Supplementary Figure 4: Correlation of node cardinality (i.e., the number of occurrences of the respective OTU in all bioindicator lists) and the node's *(a)* degree, *(b)* closeness centrality, *(c)* eigenvector centrality, *(d)* page rank, and *(e)* authority score in the bioindicator network.
